# Mechanosensitive nuclear asymmetries define a bipolar spindle scaffold to ensure mitotic fidelity

**DOI:** 10.1101/526939

**Authors:** Vanessa Nunes, Margarida Dantas, Domingos Castro, Elisa Vitiello, Irène Wang, Nicolas Carpi, Martial Balland, Matthieu Piel, Paulo Aguiar, Helder Maiato, Jorge G. Ferreira

**Author notes:** Correspondence to: Jorge G. Ferreira –, Address: Rua Alfredo Allen, 4200-135 Porto, Portugal.

## Abstract

During prophase, centrosomes need to separate and position to correctly assemble the mitotic spindle. This process occurs through the action of molecular motors, cytoskeletal networks and the nucleus. How the combined activity of these different components is spatiotemporally regulated to ensure efficient spindle assembly remains unclear. Here we show that during prophase the centrosomes-nucleus axis reorients, so that centrosomes are positioned on the shortest nuclear axis at nuclear envelope (NE) breakdown. This centrosomes-nucleus configuration depends on mechanical cues generated by mitotic chromosome condensation on the prophase nucleus. We further show these mechanosensitive cues act through SUN1/2 and NudE+NudEL to enable the polarized loading of Dynein on the NE. Finally, we observe this centrosome configuration favors the establishment of an initial bipolar spindle scaffold, facilitating chromosome capture and accurate segregation, without compromising division plane orientation. We propose that chromosome segregation fidelity depends on the mechanical properties of the prophase nucleus that facilitate spindle assembly by regulating NE-Dynein localization.

## Introduction

Chromosome segregation requires the assembly of a bipolar mitotic spindle. While multiple pathways contribute to spindle assembly [1], in human somatic cells centrosomes play a dominant role. During prophase, centrosome separation occurs independently of nuclear envelope breakdown (NEB), in a kinesin-5-dependent manner [2]. Accordingly, depletion or inhibition of kinesin-5 prevents centrosome separation, generating monopolar spindles and mitotic arrest [3]. Other players involved in the process include motors such as Dynein, both at the nucleus [4–8] and at the cell cortex [4, 9], MyosinII [10], but also actin [11] and microtubule pushing forces [9, 12]. How the forces generated by these components are functionally coordinated to ensure efficient spindle assembly remains unclear.

During prophase, centrosomes are tethered to the surface of the nuclear envelope (NE) in a Dynein-dependent manner [7, 8, 13]. Loading of Dynein occurs through multiple pathways which are under the regulation of CDK1 [7] and involve direct binding to nucleoporins [8, 13] or interaction with the LINC complex [14, 15]. Additional mechanisms which affect NE Dynein activity, but not loading [5], are also involved in centrosome separation. Importantly, by tethering centrosomes to the NE, Dynein is essential for early spindle assembly [8]. Taken together, these reports highlight the contribution of an internal signal on the prophase nucleus for early spindle assembly.

In metaphase, cortical force generators determine spindle orientation [16–19]. These are activated by external cues [20] and generate pulling forces on astral microtubules [21–23] to align the spindle with the long cell axis [24], ultimately defining the division plane. However, whether centrosome separation and early spindle assembly follow the same cortical cues remains unknown. Defining how external and internal signals are integrated during early mitosis to ensure efficient spindle assembly and robust division plane orientation is relevant, since prophase centrosome positioning is essential for accurate chromosome segregation [25, 26].

Here, we performed a high-resolution analysis of centrosome behavior during mitotic entry in human cells, followed by 3D cellular reconstruction and centrosome tracking. We show that during mitotic entry, the centrosomes-nucleus axis reorients so that centrosomes are positioned on the shortest nuclear axis. In addition, we identify a mechanosensitive nuclear signal that depends on the chromatin condensation state and enables Dynein loading on the NE. This ensures centrosome-NE tethering and correct positioning on the shortest nuclear axis. As a result, the formation of an initial bipolar spindle scaffold is facilitated, ensuring maximum exposure of kinetochores to microtubules and improving chromosome segregation fidelity. Thus, our work unveils how cytoskeletal and nuclear events are coordinated at the G2-M transition.

## Results

### Centrosomes position on the shortest nuclear axis at nuclear envelope breakdown

To characterize mitotic spindle assembly at high spatiotemporal resolution, we performed 4D imaging in HeLa cells. We observed that when cells are seeded on a substrate that does not activate integrin signaling (poly-L-lysine; PLL), centrosomes separate independently of NEB (Fig. S1A), as reported previously [2, 25]. However, when seeded on integrin-activating fibronectin (FBN), ∼82% of the cells separate their centrosomes to opposite sides of the nucleus before NEB. Moreover, cells that have an increased spreading area at NEB show longer inter-centrosome distances (Fig. S1B), suggesting that centrosome separation prior to NEB is a function of the adhesion area. To normalize cell area and shape in 2D, we seeded cells on defined FBN micropatterns and monitored centrosome dynamics, cell membrane and nuclear shape (Fig. 1A), which were subsequently reconstructed using specifically developed computational algorithms (Fig. S2). Centrosome dynamics relative to the micropattern was defined by two angles theta and phi, reflecting movements in xy (azimuth) and xz (inclination), respectively (Fig. 1B). These vary between 0° (aligned with the long axis of the pattern) and 90° (perpendicular to the pattern). We anticipated that separated centrosomes should align with the long axis of the micropattern, due to the distribution of retraction fibers imposed by extracellular matrix organization [17, 20]. However, during mitotic entry, centrosomes deviated from the underlying micropattern, as observed by the high variability of theta and phi (Fig. 1B). This was accompanied by a rotation of the nucleus relative to the long axis of the pattern, as well as a decrease in cell area (Fig. 1C). Due to the shape asymmetry of the line micropattern, we could calculate cell membrane eccentricity, which varies between 1 (completely elongated cell) and 0 (spherical cell). As cell progressed towards NEB, membrane eccentricity decrease (Fig. 1D) due to a retraction of the long cell axis (Fig. 1E, 0°) and a simultaneous increase in cell width, perpendicularly to the pattern (Fig. 1E, F; 90°; *** p<0.001). Interestingly, during the rounding process, the centrosomes and nucleus re-orientated so that centrosomes were positioned on the shortest nuclear axis at NEB (∼80% of cells; Fig. 1G-I, Movie S1).

**Figure 1:**
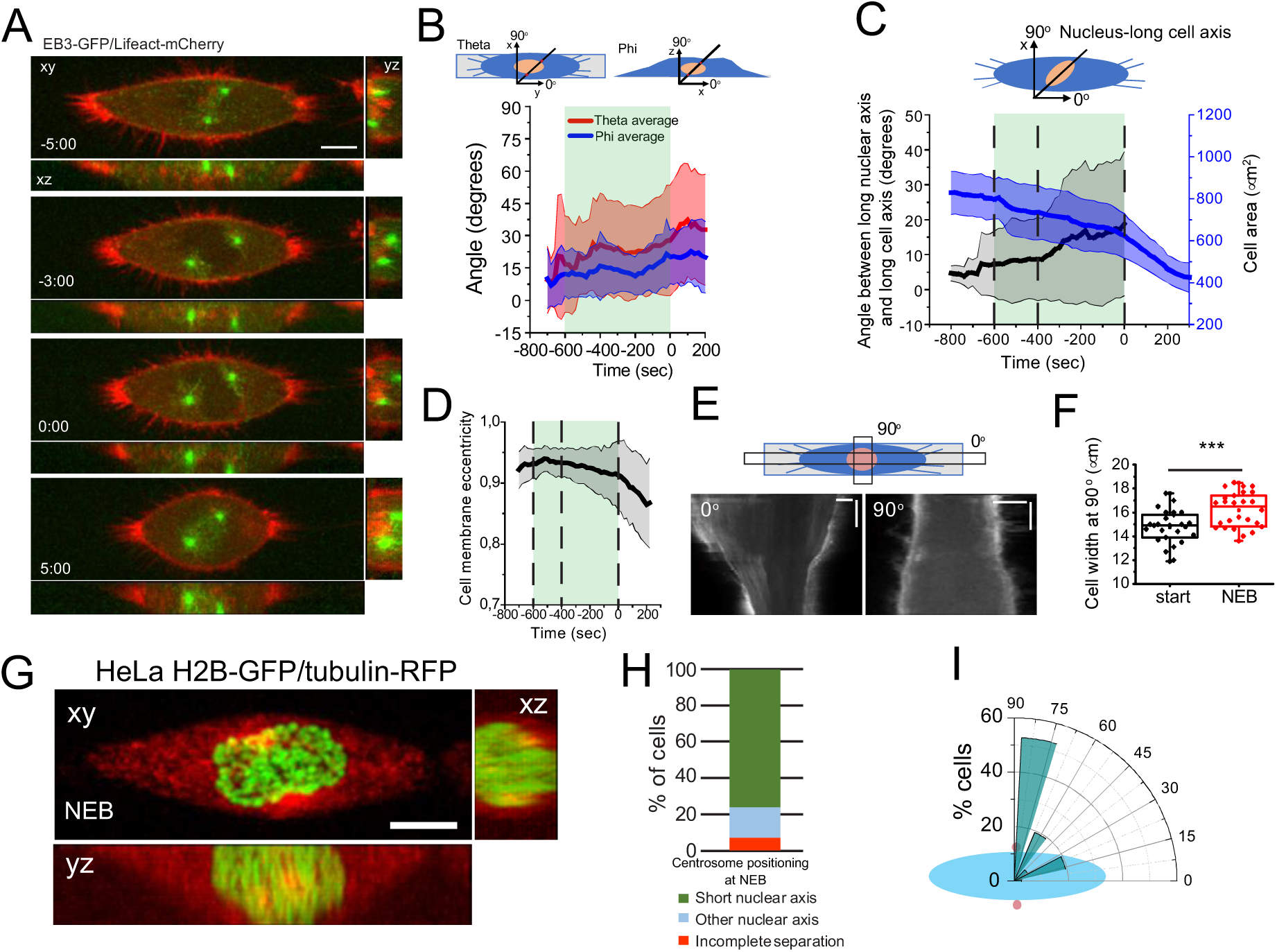
Characterization of early spindle assembly. (A) Frames from movie of a cell seeded on a line micropattern, showing movement of the centrosomes towards the shortest nuclear axis. Time is in min:sec. Time zero corresponds to NEB. (B) Characterization of centrosome orientation vector in xy (theta; red) and z (phi; blue) for cells seeded on line micropatterns (n=30). Line corresponds to average and shaded area to SD. (C) Quantification of cell area (μm^2^; blue) and angle between nucleus-long cell axis (black) for cells on line micropatterns (n=37). Lines correspond to average and shaded areas represent SD. (D) Cell membrane eccentricity of during mitotic entry for cells on line micropatterns. Line represents average value and shaded area represents SD. (E) Kymograph from cell expressing Lifeact-mCherry seeded on a line micropattern, during mitotic cell rounding. Zero degrees corresponds to the long cell axis and 90 degrees to the perpendicular orientation. (F) Cell width (μm) perpendicular to the pattern (n=16; *** p<0.001). (G) Representative frame from a movie of a cell expressing H2B-GFP/tubulin-RFP showing centrosome and nucleus orientation at NEB. (H) Quantification of centrosome separation behavior at NEB for cells seeded on line micropatterns. (I) Polar plot quantifying centrosome positioning (red circles) relative to the longest nuclear axis (blue ellipse) at NEB for cells seeded on line micropatterns. All experiments were replicated at least three times.

### Centrosome positioning requires nuclear and centrosome movement

Our observations suggest that prophase centrosome positioning on the shortest nuclear axis is a result of the combined motion of centrosomes and the nucleus. To confirm this, we analyzed the relative contribution of each component for the positioning of centrosomes on the shortest nuclear axis (Fig. 2A). We reasoned that if positioning on the shortest nuclear axis depended exclusively on centrosome movement (centrosome dominant), the centrosomes-long nuclear axis angle would tend to 90°, and the nucleus would remain aligned with the micropattern. On the other hand, if this mechanism required only nuclear rotation, then the nucleus long axis-long cell axis angle would tend to 90°, and the centrosomes movement would be residual. If both components were involved, then we would observe combined motion of both nucleus and centrosomes (Fig. 2A-E). Accordingly, when cell rounding is limited, the nucleus is aligned with the long cell axis and centrosomes deviate from the pattern (centrosome dominant; Fig. 2B, C). When cell rounding is more pronounced, the nucleus tends to rotate away from the long cell axis (nucleus dominant; Fig. 2B, D). In intermediate cases, both centrosomes and nucleus deviate from the long cell axis (Fig. 2B, E). Therefore, at earlier time points (600sec before NEB) when cells have not rounded up significantly, nuclear rotation is limited and centrosome positioning depends exclusively on centrosome motion (Fig. 2F, I). However, as rounding progresses, the contribution of nuclear rotation now plays a significant role in determining centrosomes-nuclear axis (Fig. 2G, H, J, K; *p<0.05). Overall, these observations suggest that cell rounding enables nuclear movement to facilitate the reorientation of the centrosomes-nucleus axis, so that centrosomes position on the shortest nuclear axis at NEB (Fig. 2L).

**Figure 2:**
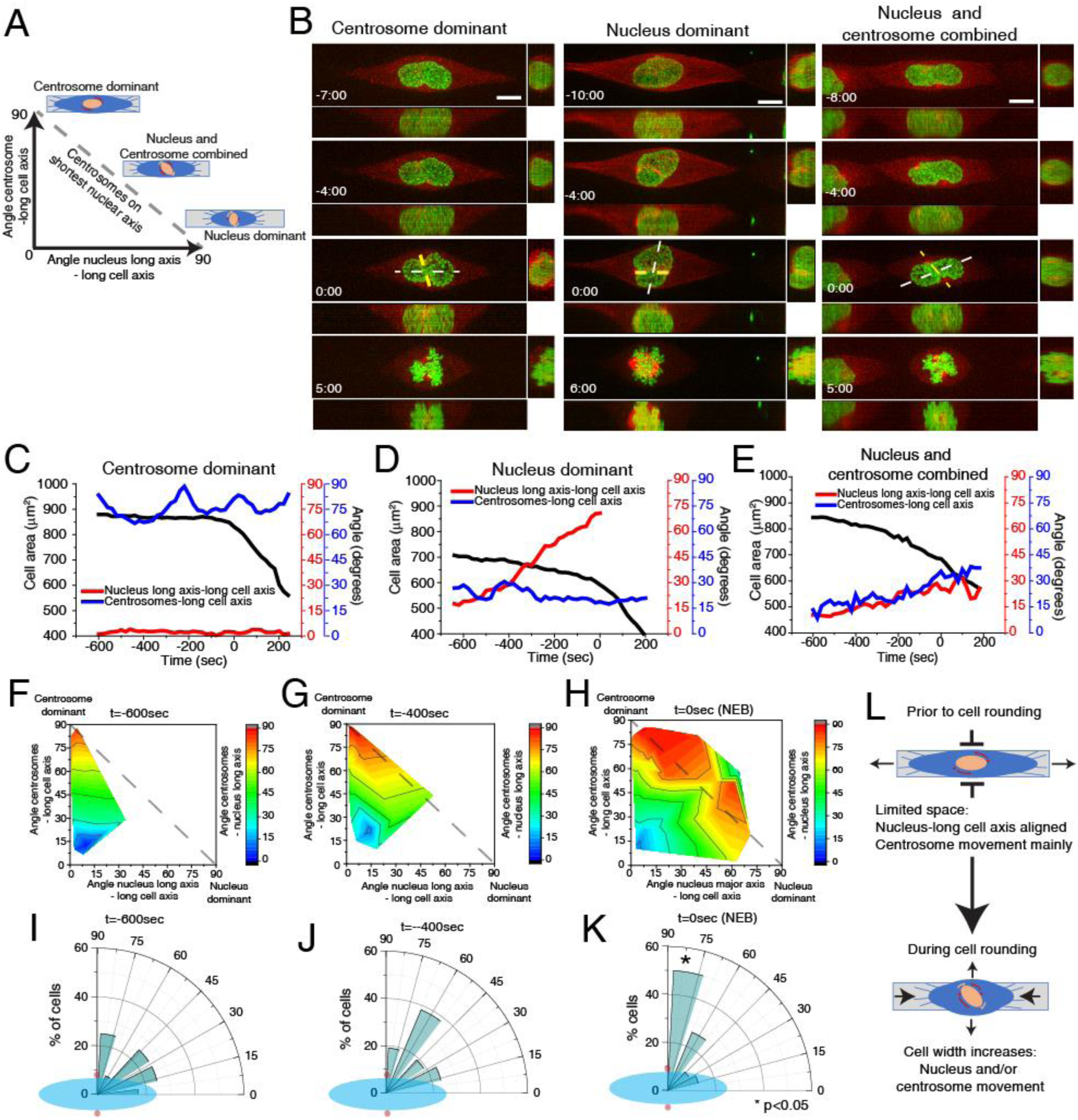
Centrosome positioning requires centrosome and nucleus movement. (A) Positioning of centrosomes on the shortest nuclear axis can be achieved in three different ways: centrosome movement, nucleus movement, or both. (B) Selected frames from movies of HeLa cells expressing H2B-GFP/alpha-tubulin-RFP seeded on line micropatterns to show the three positioning modes (n=38). White line shows the long nuclear axis and yellow lines show centrosomes axis. Time lapse is 20 sec. Time is in min:sec. Time zero corresponds to NEB. Scale bars, 10 μm. Representative plots showing the correlation between centrosome-long cell axis (blue), long nuclear axis-long cell axis (red) and cell area (black) for centrosome dominant (C), nucleus dominant (D) and nucleus-centrosome combined (E) behaviors. Quantification of the contribution of centrosome displacement (angle between centrosomes-long cell axis) and nucleus displacement (angle nucleus long axis-long cell axis) for centrosome positioning on the shortest nuclear axis (angle centrosomes-long nuclear axis) at −600sec (F), −400 sec (G) and NEB (H). Distribution of centrosome positioning (red circles) relative to the longest nuclear axis (blue ellipse) at −600sec (I), −400sec (J) and NEB (K). (L) Before cell rounding, centrosome-nucleus axis orientation depends mainly on centrosome movement due to the limitation in space. During mitotic rounding, cell width increases, allowing nuclear rotation.

### Centrosome positioning on the shortest nuclear axis depends on cell adhesion area but not cell shape

Our initial observations were obtained with cells seeded on line micropatterns that have a highly polarized shape. To determine whether centrosome positioning on the shortest nuclear axis was a result of shape polarization or a more general feature, we followed cells on large circles, small circles and rectangles during mitotic entry (Fig. 3A). Strikingly, changing from a polarized shape such as a rectangle to an unpolarized large circle did not block the capacity of centrosomes to position on the shortest nuclear axis (78% of cells on rectangles and 75% of cells on large circles; Fig. 3A-D). However, correct positioning depended on the initial spreading area, as seeding cells in small circles led to erratic centrosome movement and only 25% of cells placed centrosomes on the short nuclear axis (Fig. 3A, B, E; *p=0.02). In addition, cell shape did not interfere with nucleus orientation relative to the micropattern (Fig. 3F-H), although nuclear shape did change when cells were seeded on small circles (Fig. 3I; ***p<0.001), with nuclei becoming more rounded. As a result, cells on small circles lost the coordination of movement between centrosomes and nucleus, when compared to the other micropatterns. Taken together, these data indicate that centrosomes position on the shortest nuclear axis at NEB in a cell area-dependent manner but independently of cell shape.

**Figure 3:**
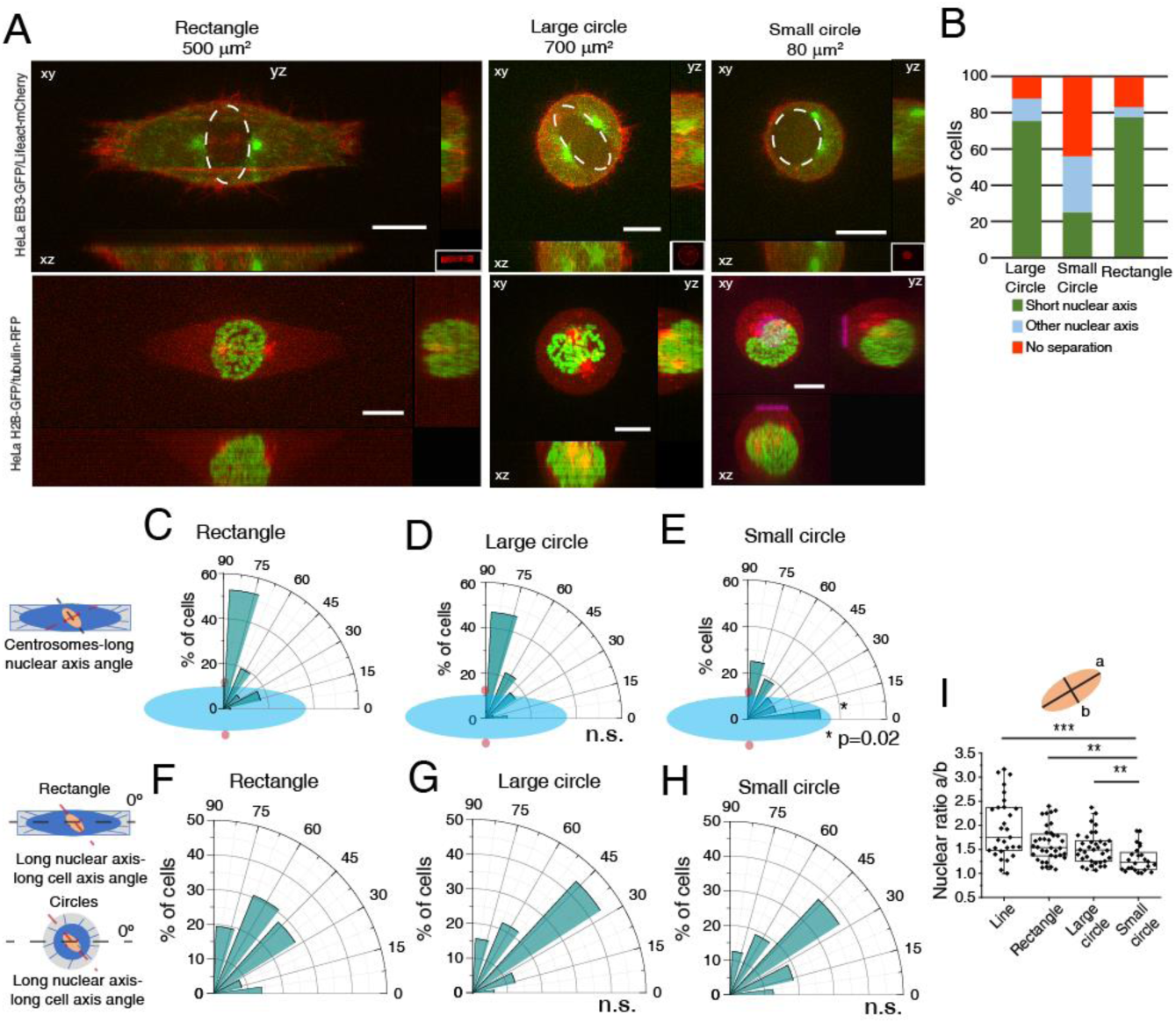
Centrosome positioning on the shortest nuclear axis does not depend on shape polarization. (A) Representative images from cells at NEB, expressing EB3-GFP/Lifeact-mCherry (top panels) or H2B-GFP/alpha-tubulin-RFP (bottom panels) seeded on rectangles (500μm^2^; n=36), large circles (700μm^2^; n=32) or small circles (80μm^2^; n=16), showing lateral projections (xz and yz). Scale bars, 10μm. Ellipses highlight nuclear shape at NEB. (B) Quantification of centrosome separation behavior at NEB for cells seeded on the different micropatterns. Polar plot quantifying centrosome positioning (red circles) relative to the longest nuclear axis (blue ellipse) at NEB for cells seeded on rectangles (C), large circles (D) or small circles (E). Polar plot quantifying alignment of the long nuclear axis with the long cell axis at NEB for cells on rectangles (F), large circles (G) and small circles (H). For cells seeded on circles, zero was defined horizontally. (I) Quantification of nuclear shape asymmetry, as defined by the ratio long nuclear axis/short nuclear axis, for cells seeded on different micropatterns (***p<0.001; **p<0.01).

### Cell rounding allows the centrosomes-nucleus axis to reorient in prophase

Metaphase spindle orientation is determined by the distribution of actin-based retraction fibers and this depends on extracellular matrix organization [17, 20]. Therefore, it would be reasonable to assume that centrosomes should orient according to the same cues during prophase. However, our results show that prior to NEB (and simultaneously with cell rounding), the centrosomes-nucleus axis reorients away from the underlying retraction fiber distribution imposed by the micropattern. This suggests that the rounding process changes the manner in which cells interact with the extracellular matrix. To confirm this, we performed Traction Force Microscopy (TFM) analysis on cells seeded on rectangles (Fig. S3A). Under these conditions, cells showed a well-defined traction axis that correlated with the initial centrosome separation axis (theta; Fig. S3A-C). Upon mitotic rounding, both cell area and the contractile energy exerted on the substrate decreased (Fig. S3D), leading us to conclude that mitotic rounding decreases the force exerted by the cell on the substrate. These observations, together with our previous results, suggest that blocking cell rounding could affect centrosome positioning. To test this we decided to express a mutant form of Rap1 (Rap1Q63E; Rap1*) that interferes with focal adhesion disassembly, effectively blocking mitotic rounding [27]. Accordingly, Rap1* expression affected cell rounding, when compared to controls (Fig. 4A, B, D, E). Centrosome movement was also affected in Rap1* cells, as theta and phi were decreased when compared to controls (Fig. 4B, E; ***p<0.001). Importantly, the inability of Rap1* cells to round up led to a significant impairment in nuclear rotation when compared to controls (Fig. 4C, F; *p<0.05). This eventually resulted in a failure to position centrosomes on the shortest nuclear axis at NEB (Fig. 4G-J; **p<0.01). We conclude that cell rounding is required to establish the centrosomes-nucleus axis during prophase.

**Figure 4:**
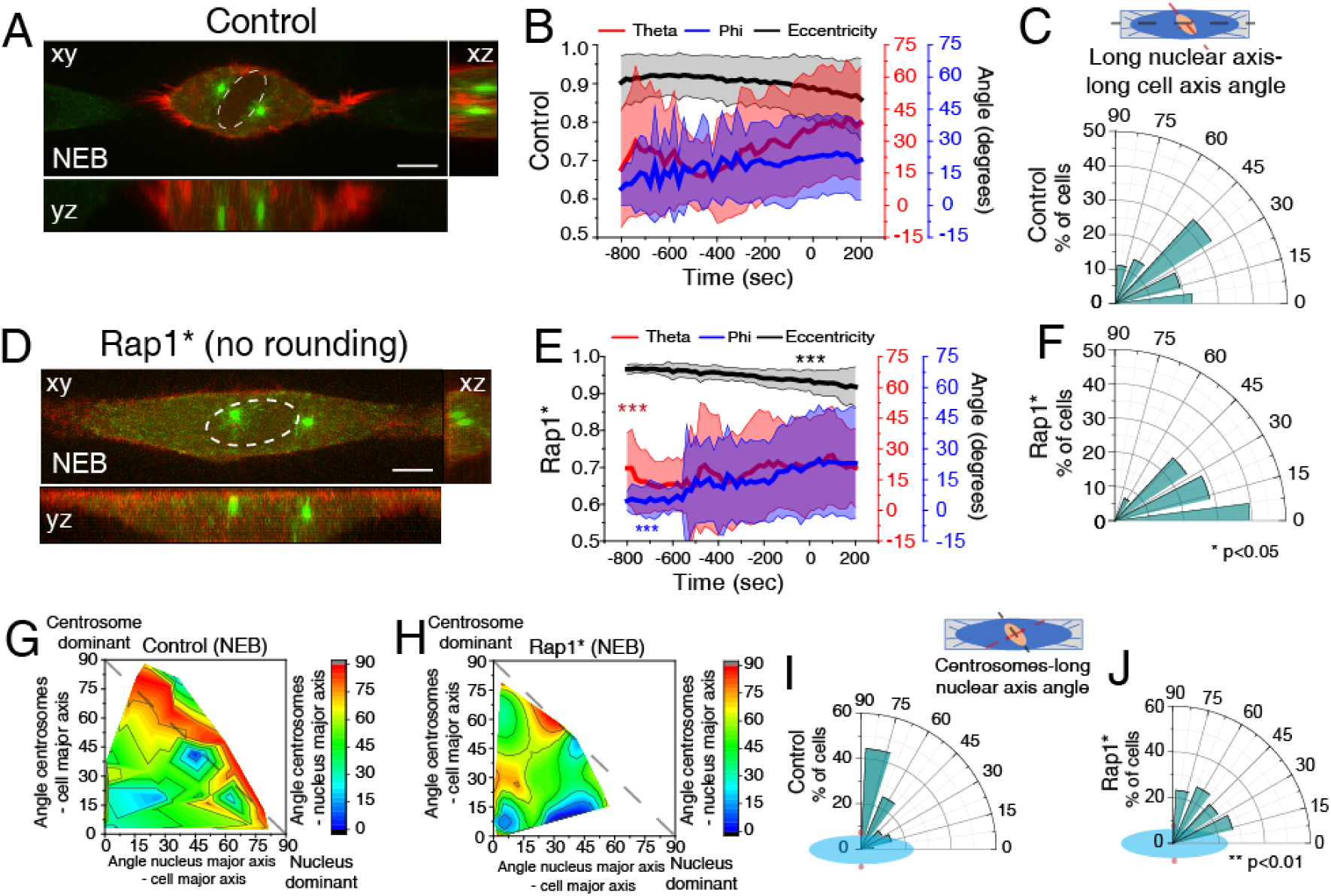
Cell rounding is required to establish the centrosomes-nucleus axis at NEB. Representative time-frame from movie of HeLa cell expressing EB3-GFP/Lifeact-mCherry treated with DMSO (A; n=38) or Rap1* (D; n=28) at NEB. Correlation of cell membrane eccentricity (black), theta (red) and phi (blue) for controls (B) and Rap1* (E). Lines correspond to average values and shaded areas correspond to SD. Inhibiting cell rounding limits centrosome movement (*** p<0.001). Polar plot quantifying alignment of the long nuclear axis with the long cell axis at NEB for DMSO (C) or Rap1* (F; * p<0.05). Quantification of the contribution of centrosome displacement (angle between centrosomes-long cell axis) and nucleus displacement (angle nucleus long axis-long cell axis) for centrosome positioning on the shortest nuclear axis (angle centrosomes-long nuclear axis) for controls (G) and Rap1* (H) at NEB. Polar plot quantifying centrosome positioning (red circles) relative to the longest nuclear axis (blue ellipse) at NEB for controls (I) and Rap1* (J; ** p<0.01). All experiments were replicated at least three times.

### Dynein on the nuclear envelope is required for centrosome-nucleus reorientation during prophase

Next, we wanted to determine which factors influence the positioning of centrosomes on the shortest nuclear axis. It is well known that kinesin-5 is essential for centrosome separation [2, 3]. To assess whether it is also required to position centrosomes on the shortest nuclear axis, we treated cells with an Eg5 inhibitor (STLC), when centrosomes were already on opposite sides of the nucleus (Late stage) or when centrosomes were still not fully separated (Early stage). Early Eg5 inhibition significantly decreased inter-centrosome distance, preventing centrosome positioning on the shortest nuclear axis (Fig. S4A-C). When Eg5 was inhibited in the Late stage, centrosomes moved towards the shortest nuclear axis, simultaneously with mitotic cell rounding (Fig. S4D, dashed line). We concluded that kinesin-5 is required for initial centrosome separation but not directionality.

Centrosome positioning on the shortest nuclear axis at NEB could rely on signals coming from the cytoskeleton, centrosomes or intrinsic nuclear cues. To test this, we experimentally uncoupled all components. Centrosome-nucleus tethering in prophase requires Dynein loading on the NE, which occurs by two pathways involving RanBP2-BicD2 and Nup133-CENP-F-NudE/NudEL [7, 8, 13, 28]. Interestingly, Dynein is also involved in centrosome separation [6, 29], making it a likely candidate to mediate centrosomes-nucleus orientation. Accordingly, depletion of either NudE+NudEL or BicD2 led to centrosome detachment from the NE (Fig. 5A; Movie S2). Consequently, centrosomes no longer positioned on the shortest nuclear axis (Fig. 5B-D; ***p<0.001). Moreover, both depletions impaired rotation of the nucleus relative to the long cell axis (5E-G; ***p<0.001). Unexpectedly, cell rounding was differentially affected by the two depletions (Fig. 5H; ***p<0.001), suggesting that BicD2 is required for nuclear rotation independently of cell rounding. Depletion of total Dynein Heavy Chain (DHC) induced similar centrosome positioning and nuclear rotation defects, which could be due to a delayed cell rounding (Fig. S5A-C; ***p<0.001). We conclude that centrosome-nucleus coupling through NE Dynein is essential for centrosome-nucleus axis orientation at NEB.

**Figure 5:**
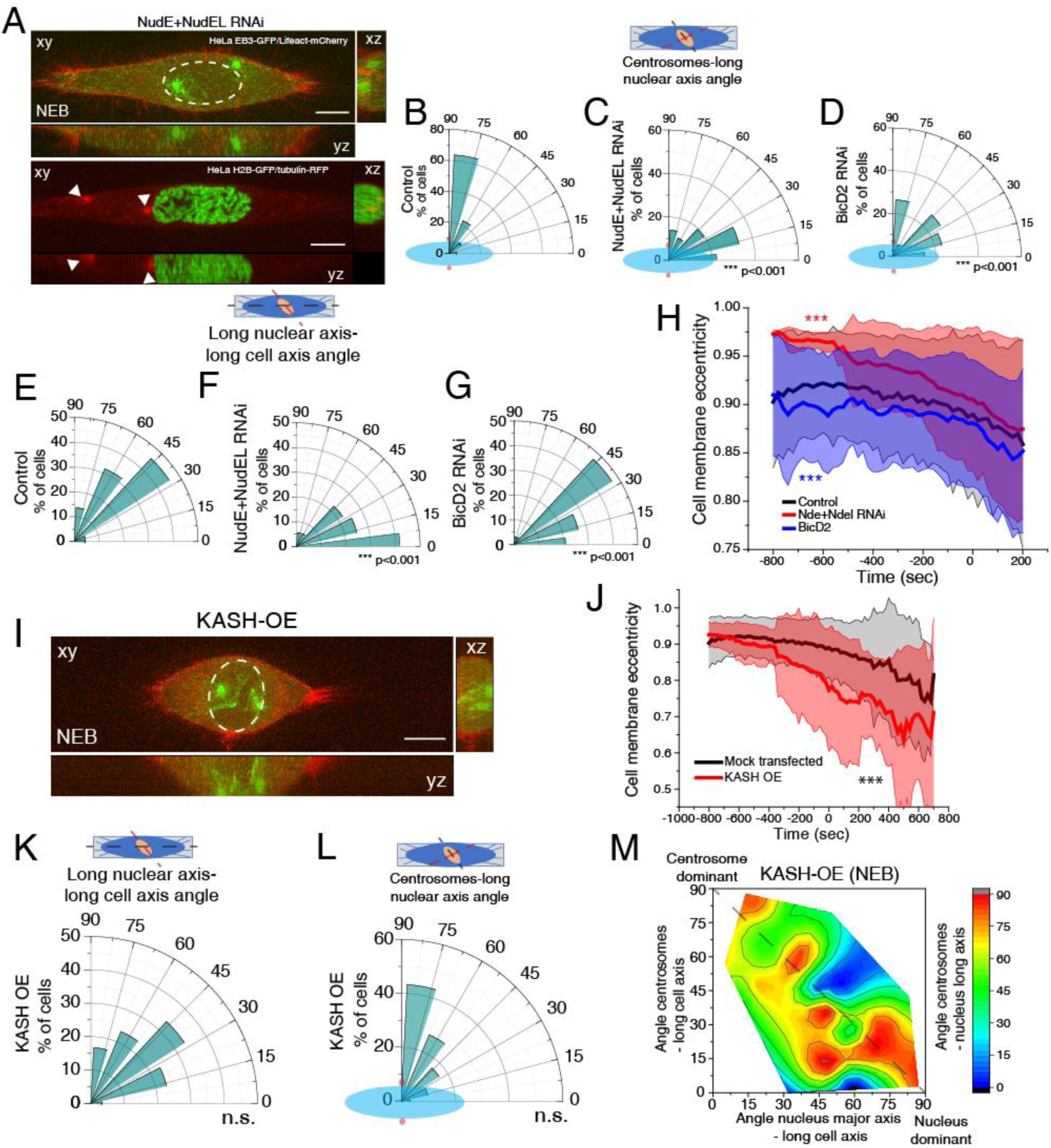
Dynein on the nuclear envelope is required to establish the centrosomes-nucleus axis at NEB. (A) Images of HeLa cells expressing EB3-GFP/Lifeact-mCherry (top panel; n=36) or H2B-GFP/tubulin-RFP (bottom panel; n=25) treated with NudE+NudEL RNAi. White arrowheads indicate centrosome position. Time-lapse is 20 sec. Scale bar, 10μm. Polar plot quantifying centrosome positioning (red circles) relative to the longest nuclear axis (blue ellipse) at NEB for controls (B), cells treated with RNAi for NudE+NudEL (C) or BicD2 (D; n=34). Polar plot quantifying alignment of the long nuclear axis with the long cell axis at NEB for controls (E), NudE+NudEL RNAi (F; *** p<0.001) or BicD2 RNAi (G; *** p<0.001). (H) Cell membrane eccentricity for controls, NudE+NudEL RNAi and BicD2 RNAi. Lines correspond to average values and shaded areas correspond to SD (*** p<0.001). (I) Time-frame from cell expressing EB3-GFP/Lifeact-mCherry, transfected with a KASH-GFP construct (n=30). Scale bar, 10μm. (J) Cell membrane eccentricity controls and KASH-GFP transfected cells. Lines correspond to average values and shaded areas correspond to SD (***p<0.001). (K) Polar plot quantifying alignment of the long nuclear axis with the long cell axis at NEB for cells expressing KASH-GFP. (L) Polar plot quantifying centrosome positioning (red circles) relative to the longest nuclear (blue ellipse) at NEB for cells expressing KASH-GFP. (M) Quantification of the contribution of centrosome displacement (angle between centrosomes-long cell axis) and nucleus displacement (angle nucleus long axis-long cell axis) for centrosome positioning on the shortest nuclear axis (angle centrosomes-long nuclear axis) at NEB for cells expressing KASH-GFP. All experiments were replicated at least three times.

Dynein can be found in different subcellular localizations. During later stages of mitosis, it localizes to the cell cortex through the LGN-Gαi-NuMA complex, and this can be prevented by inhibiting Gαi activity with pertussis toxin (PTx) [19, 30]. To determine whether cortical Dynein is also involved in centrosome-nucleus reorientation, we treated cells with PTx during mitotic entry (Fig. S5D). Under these conditions, cells rounded up prematurely (Fig. S5E; ***p<0.001) and the nucleus showed increased rotation (Fig. S5F; **p=0.002). Overall, this did not prevent centrosomes positioning on the shortest nuclear axis at NEB (Fig. S5G), since these cells are still able to load Dynein on the NE (Fig. S5H, I). Taken together, our results indicate that cortical Dynein is not required for centrosomes-nucleus reorientation during prophase.

Finally, we tested whether *Linker of nucleoskeleton and cytoskeleton* (LINC)-mediated nucleus-cytoskeletal coupling is necessary for centrosome positioning, as this complex is required for NE Dynein loading and centrosome tethering during neuronal development [14, 15]. We expressed a KASH-GFP construct (Fig. 5I), which displaces endogenous Nesprins from the NE, disrupting the LINC complex [31, 32] and consequently its connection with the cytoskeleton. Under these conditions, cell rounding was faster than in controls (Fig. 5J; ***p<0.001) but nuclear rotation was not affected (Fig. 5K). Consequently, centrosomes still positioned on the shortest nuclear axis at NEB (Fig. 5L), although the centrosome-nucleus reorientation shifted towards a nucleus-dependent mode (Fig. 5M). Overall, we concluded that centrosomes need to be attached to the surface of the NE in a Dynein-dependent manner to position on the shortest nuclear axis at NEB.

### Chromatin condensation is required for centrosome positioning

We showed that prophase centrosome-nucleus coupling mediated by Dynein is required for proper centrosome positioning. However, this does not explain the bias towards the shortest nuclear axis. We reasoned there must be some intrinsic nuclear property that could provide the cues for centrosome positioning. We started by analyzing DHC-GFP distribution on the NE during prophase with both live-cell imaging (Fig. 6A, Movie S3) and immunofluorescence analysis (Fig. 6B) and found that Dynein was distributed in an asymmetric manner on the NE, concentrating on curved areas of the envelope (Fig. 6A, B, D). To further characterize NE behavior, we measured active NE fluctuations in cells expressing the nucleoporin POM121-3xGFP. We calculated deviations from the median and the maximum amplitude for each NE coordinate and time point. Fluctuations were analysed on the regions corresponding to the long (green triangles) and short (red triangles) nuclear axes (Fig. 6E) and revealed significant differences between the long and short nuclear axes (Fig.6 F, G). Overall, these data indicate that the prophase NE has spatial asymmetries.

**Figure 6:**
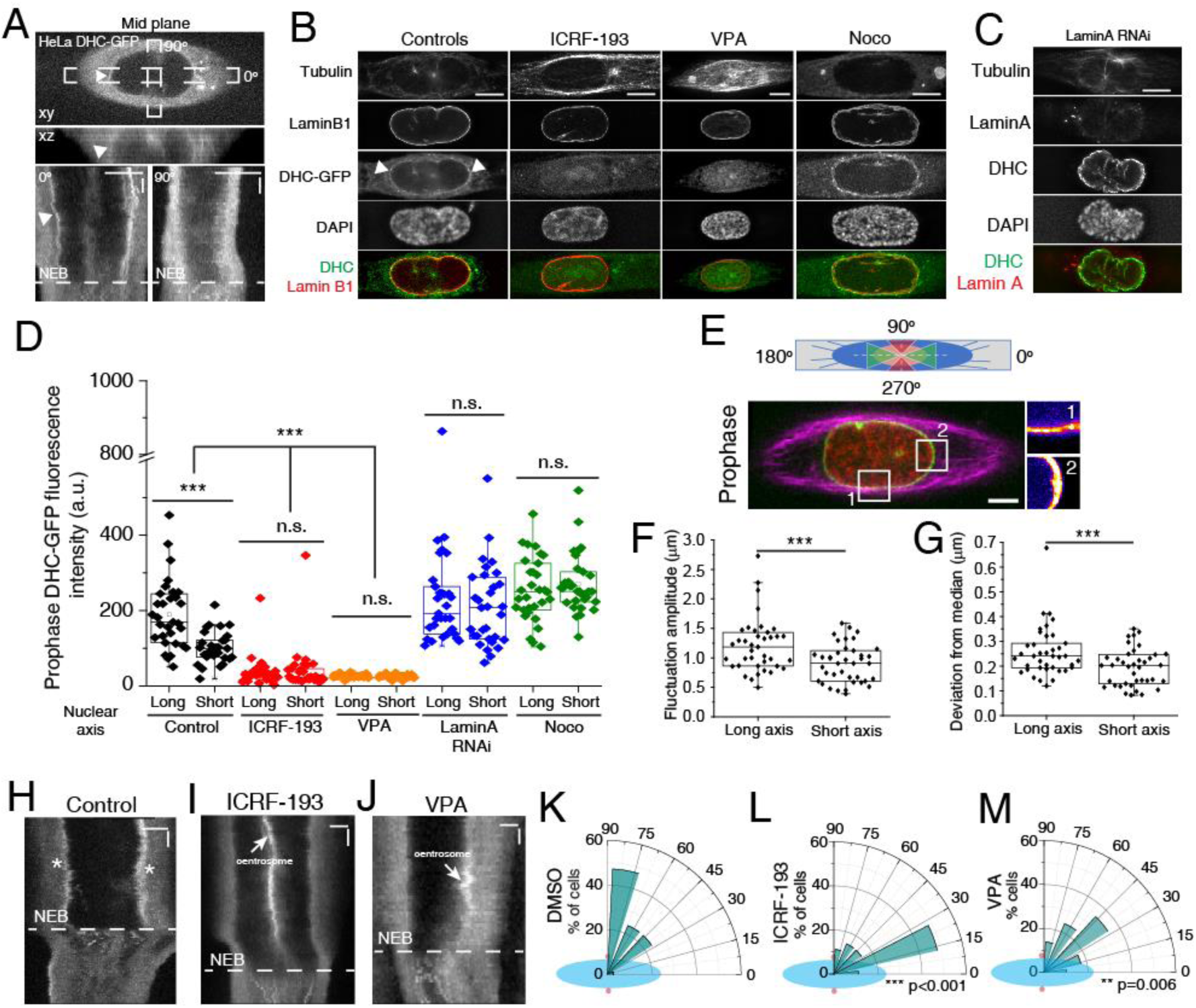
Spatial asymmetries of the prophase nucleus determine centrosome positioning. (A) Kymograph of cell expressing DHC-GFP (white arrowheads indicate polarized NE-Dynein). Horizontal scale bar, 10μm. Vertical scale bar 100sec. (B) Immunofluorescence images of DHC-GFP, Lamin B1, alpha-tubulin and DAPI for controls, ICRF-193, VPA and Noco. Images correspond to deconvolved intensity sum projections. White arrowheads indicate asymmetric localization of DHC-GFP. Scale bar, 10μm. (C) Immunofluorescence images of DHC, Lamin A, alpha-tubulin and DAPI for cells treated with Lamin A RNAi. (D) Quantification of NE-associated DHC-GFP fluorescence intensity in prophase for controls (n=31), ICRF-193 (n=30), VPA (n=40), Lamin A RNAi (n=31) and Noco (n=31) (*** p<0.001; n.s. – not significant). For quantification purposes, sum-projected images covering the nucleus region were used. (E) Regions analyzed for nuclear fluctuations in cells expressing POM121-3xGFP/H2B-mCherry/SiR-tubulin. Scale bar, 10μm. Insets represent maximum temporal projections of POM121. Long nuclear axis corresponds to green triangles and short nuclear axis corresponds to red triangles. Fluctuation amplitude (F) and deviation from the median (G) for nuclear envelope fluctuations of prophase cells (n=39; *** p<0.001). Kymographs from control (H; n=41), ICRF-193 (I; n=13) and VPA-treated cells (J; n=8) expressing DHC-GFP. Asterisks indicate NE-Dynein in control cells. Dynein does not accumulate on the NE with ICRF-193 or VPA treatment. Horizontal scale bar, 10μm. Vertical scale bar 100sec. Polar plot quantifying centrosome positioning (red circles) relative to the longest nuclear axis (blue ellipse) at NEB for cells treated with DMSO (K; n=39), ICRF-193 (L; n=27; ***p<0.001) and VPA (M; n=29; ***p<0.001). All experiments were replicated at least three times.

During interphase, Lamin A and chromatin are the main regulators of the nuclear mechanical response [33, 34], that mediate the response to large and small deformations, respectively. To test their relative contribution for the nuclear asymmetry, we analyzed Dynein distribution on the NE following depletion of Lamin A by RNAi or after treatment with ICRF-193, a Topoisomerase II (TopoII) inhibitor, or Valproic acid (VPA), a histone deacetylase inhibitor (HDAC). Treatment with either TopoII or HDAC inhibitors should interfere with chromatin structure, decreasing overall nuclear stiffness [35]. Interestingly, inhibition of either TopoII or HDACs induced a loss of Dynein specifically on the NE (Fig. 6B, D, H-J), without interfering with Dynein recruitment to the kinetochore or cell cortex (Fig. S6). Moreover, BicD2 and NudE localization also seemed unaffected (Fig. S7A, B). Overall, these data suggest that mitotic chromosome condensation during prophase is required for NE-Dynein recruitment, independently of its known nucleoporin adaptors. On the other hand, Lamin A depletion induced a loss of Dynein polarization, without affecting its recruitment to the NE (Fig. 6C, D). This was similar to what we observed following incubation with Nocodazole (Noco; Fig. 6B, D) and allowed us to conclude that an intact microtubule network and nuclear lamina are required to restrict the spatial distribution of NE-Dynein. Moreover, the structure of the nuclear lamina as visualized by LaminB1 immunostaining did not seem altered after treatment with Noco, ICRF-193 or DHC RNAi (Fig. S7C).

Since modifying chromatin condensation disrupted Dynein loading, we assessed whether these treatments also affected centrosome positioning. Accordingly, treatment with ICRF-193 or VPA prevented centrosome positioning on the shortest nuclear axis, when compared to controls (Fig. 6K-M; Fig. S8A, C; ***p<0.001). However, they had different effects on nuclear rotation (Fig. S8B, D; ***p<0.001). Inversely, Lamin A depletion did not affect centrosome positioning at NEB, nuclear rotation or cell rounding, when compared to controls (Fig. S8E-H). However, instead of migrating along the NE, centrosomes often deformed the nucleus to generate a local short axis (Fig. S8E, bottom panel). Accordingly, prophase nuclei in Lamin A RNAi cells were more irregular (Fig. S8I) and positioned their centrosomes further away from the nuclear centroid (Fig. S8J). Overall, we conclude that chromatin condensation is required for centrosome positioning on the shortest nuclear axis at NEB.

### Mechanical confinement is sufficient to load Dynein on the nuclear envelope

Interphase chromatin condensation state affects nuclear stiffness and shape [34–36]. Given our observations that interfering with chromatin condensation affects Dynein loading and that compressive forces can induce a reversible chromatin condensation [37], we wondered whether mechanical confinement of the nucleus could rescue Dynein loading in cells treated with either ICRF-193 or VPA (Fig. 7A). To test this, we applied a reversible confinement on cells and analyzed DHC-GFP localization using live-cell imaging and immunofluorescence analysis. Confinement of control cells decreased DHC-GFP signal on the NE (Fig. 7B, C), possibly due to a decrease in Dynein density [5]. Confinement release was sufficient to restore NE-Dynein localization (Fig. 7B, C), albeit without restoring its polarized distribution (Fig. 7E, F). Importantly, confinement and release of ICRF-193 or VPA-treated cells, was sufficient to rescue NE-Dynein loading, without restoring polarization (Fig. 7B, D-F). We conclude that mechanical compression of the nucleus, when chromosome condensation is pharmacologically altered, is sufficient to rescue NE-Dynein loading.

**Figure 7:**
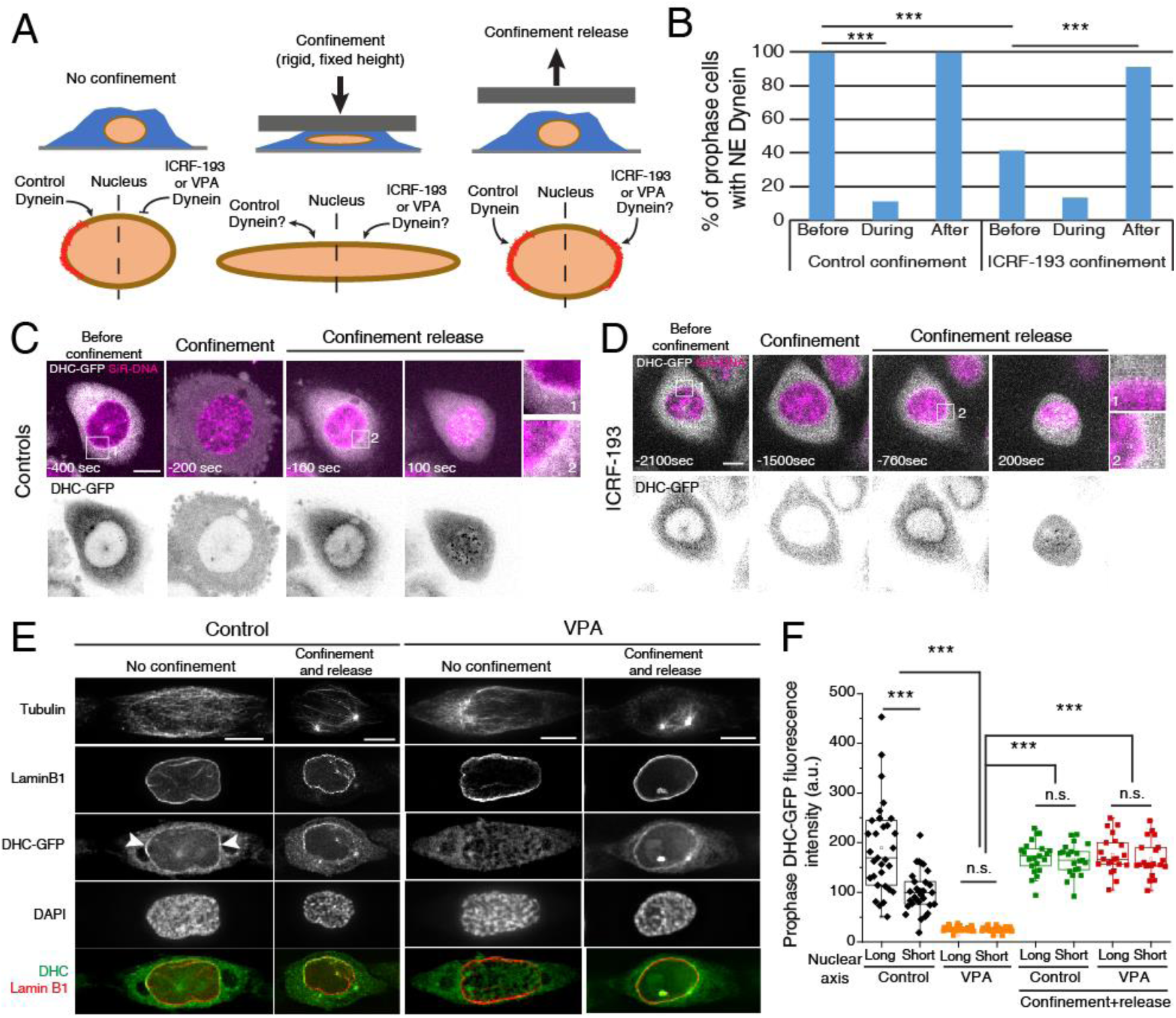
Dynein is loaded on the nuclear envelope following mechanical stimulation. (A) Experimental setup to test the effect of confinement on Dynein loading on the prophase nucleus. (B) Quantification of the percentage of prophase cells with Dynein on the NE before, during and after confinement (controls, n=14; ICRF-193, n=22; ***p<0.001). (C) Frames from movie of control HeLa cells expressing DHC-GFP and SiR-DNA (n=27) before, during and after confinement. (D) Frames from movie of HeLa cells treated with ICRF-193 (n=23), expressing DHC-GFP and stained with SiR-DNA during mechanical stimulation. (E) Immunofluorescence images of controls (n=31) and VPA-treated cells (n=41) expressing DHC-GFP, without confinement (“No confinement”) or 30 min after confinement release (“Confinement and release”) stained for tubulin, LaminB1 and DAPI. White arrowheads indicate polarized NE-Dynein in unconfined control cells. (F) Quantification of DHC-GFP intensity for controls (n=31), controls with confinement and release (n=21), VPA (n=41) and VPA with confinement and release (n=21) (*** p<0.001; n.s. – not significant). For quantification purposes, sum-projected images covering the nucleus region were used. Time lapse is 20sec. Time zero corresponds to NEB. Scale bar, 10μm. All experiments were replicated at least three times.

Next, to determine whether the mechanical regulation of Dynein recruitment to the NE required the known NE adaptors, we treated cells with ICRF-193 in combination with depletion of BicD2 or NudE/NudEL. Then, we subjected these cells to mechanical confinement and assessed NE-Dynein levels. Strikingly, mechanical confinement in the absence of NudE/NudEL was not able to rescue Dynein loading on the NE, contrary to BicD2-depleted cells (Fig. 8A-C). These observations indicate that the mechanical recruitment of Dynein to the NE occurs through the NudE/NudEL pathway. We then set out to test how the mechanical stimulus could be transmitted across the NE.

**Figure 8:**
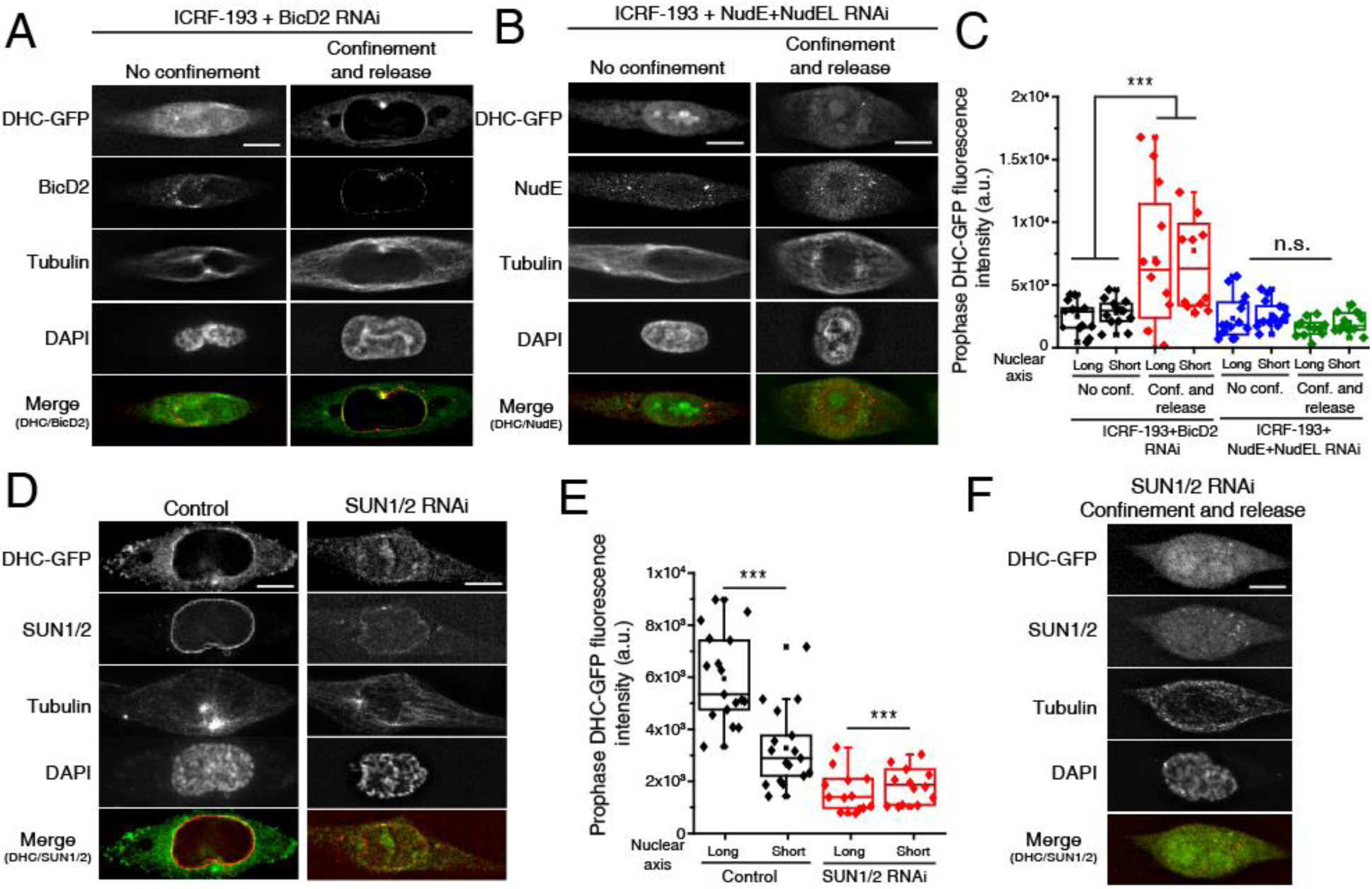
SUN1/2 and NudE+NudEL are required for the mechanosensitive loading of Dynein. (A) Immunofluorescence images of DHC-GFP, BicD2, alpha-tubulin and DAPI for a cell treated with ICRF-193 and BicD2 RNAi before and after confinement. (B) Immunofluorescence images of DHC-GFP, NudE, alpha-tubulin and DAPI for a cell treated with ICRF-193 and NudE+NudEL RNAi before and after confinement. (C) Quantification of DHC-GFP intensity for cells treated with ICRF-193+BicD2 without confinement (n=13), ICRF-193+BicD2 after confinement release (n=12), ICRF-193+NudE RNAi without confinement (n=15) and ICRF-193+NudE RNAi after confinement release (n=13). (D) Immunofluorescence images of DHC-GFP, SUN1/2, alpha-tubulin and DAPI for control cells (n=17) and cells treated with SUN1/2 shRNA (n=14). (E) Quantification of DHC-GFP intensity for controls and cells treated with SUN1/2 shRNA. (F) Immunofluorescence images of DHC-GFP, SUN1/2, alpha-tubulin and DAPI for cells treated with SUN1/2 shRNA after confinement release. Scale bars, 10μm.

**Figure 9:**
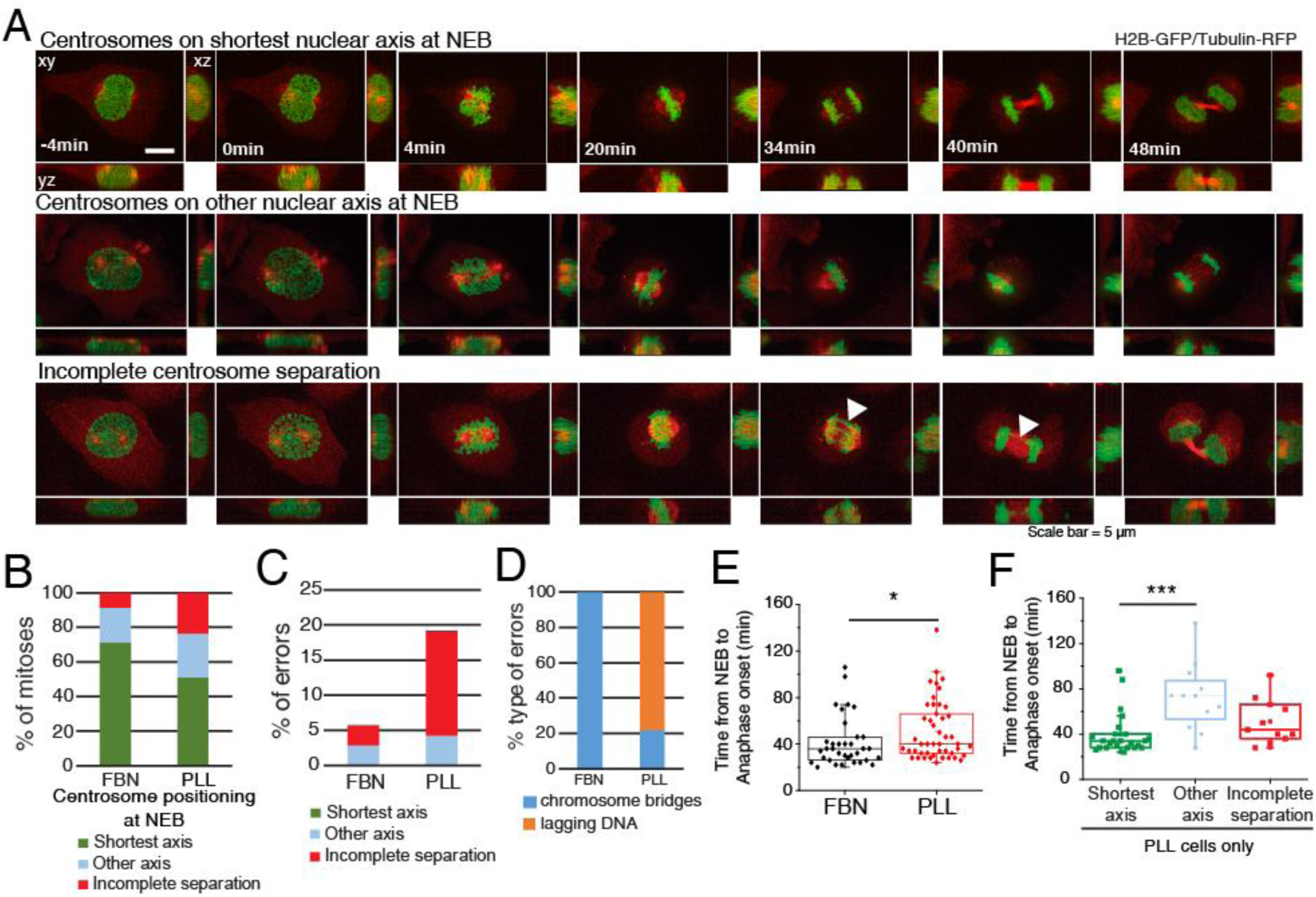
Centrosome positioning on the shortest nuclear axis facilitates spindle assembly. (A) HeLa cells expressing H2B-GFP/alpha-tubulin-RFP during mitosis. Top panel represents a cell with centrosomes on shortest nuclear axis at NEB, middle panel represents a cell with centrosomes on a random nuclear axis and bottom panel represents a cell with incomplete centrosomes separation. Time-lapse is 2 min. Scale bar, 5μm. (B) Proportion of cells that place centrosomes on shortest axis (green), other axis (blue) or have incomplete separation (red) at NEB, depending on the coating (FBN or PLL). Quantification of the proportion (C) and type of missegregation events (D) for cells seeded on FBN (n=35) or PLL (n=47). (E) Quantification of the time between NEB and anaphase onset for cells seeded on FBN or PLL (*** p<0.001). (F) Timings from NEB to anaphase onset, for cells seeded on PLL, according to their centrosome separation status (*** p<0.001; n.s. – not significant).

Transduction of mechanical forces to the nucleus requires an intact SUN-KASH complex [38]. Importantly, SUN1/2 were shown to be required for NE Dynein loading during neuronal development [14, 15] and mitotic entry [39]. To test whether SUN proteins could mediate the mechanoresponsive recruitment of Dynein to the NE during prophase, we depleted SUN1/2 using shRNA in HeLa cells expressing DHC-GFP. Upon SUN1/2 depletion, Dynein levels on the NE were significantly reduced (Fig. 8E; *** p<0.001) and could not be recovered after mechanical mechanical stimulation (Fig. 8F). Overall, our data suggests that SUN1/2 transmit mechanical forces across the NE to allow loading of Dynein via the Nup133/CENP-F/NudE-NudEL pathway.

### Centrosome positioning on the shortest nuclear axis facilitates spindle assembly

Centrosome positioning at NEB is known to affect mitotic fidelity [25, 26]. Here, we tested specifically whether positioning on the shortest nuclear axis affects spindle assembly efficiency. We imaged cells on FBN or PLL and correlated centrosome positioning, mitotic timing and missegregation events (Fig. 8A, Movies S4 and S5). Seeding cells on PLL affected centrosome separation (“incomplete separation”; 23% for PLL and 8% for FBN) and positioning on the shortest nuclear axis (51% for PLL and 72% for FBN), when compared to FBN (Fig. 8B). Consequently, PLL-seeded cells had increased missegregation events (19.2%) when compared to FBN (5.7%) (Fig. 8C). These were mainly lagging chromosomes in cells with incomplete centrosome separation (Fig. 8D), as was described previously [26]. We conclude that the extent of centrosome separation increases chromosome segregation fidelity. Next, we determined whether centrosome positioning affects mitotic timing. Cells on PLL had a significant delay in anaphase onset when compared to cells on FBN (Fig. 8E, * p<0.05). This delay was due to cells that separate, but do not position centrosomes on the shortest nuclear axis (“other axis”; 72±29 min, *** p<0.001), as opposed to cells with centrosomes on the “shortest axis” (40±18min). We conclude that centrosome positioning on the shortest nuclear axis ensures timely progression through mitosis.

## Discussion

Mitotic spindle assembly is essential for the fidelity of chromosome segregation. Accordingly, delays in centrosome separation [25, 26] or alterations in spindle geometry [40] often lead to chromosomal instability (CIN). Therefore, understanding the mechanisms regulating centrosome separation and bipolar spindle assembly is critical to determine the causes underlying CIN.

At the transition from G2 to mitosis, interphase adhesion complexes disassemble [27, 41], cell margins retract [42] and microtubule dynamics change [43, 44]. This leads to a reorganization of the cytoskeleton, which is required to form a stiff mitotic cortex [45] that facilitates bipolar spindle assembly [46]. During this stage, Dynein is loaded on the NE through multiple pathways mediated by RanBP2 [13], Nup133 [8], the LINC complex [14, 15] and the nuclear lamina [5]. Here, we show that mitotic cell rounding in combination with NE Dynein are essential to allow reorientation of the centrosomes-nucleus axis, so that centrosomes position on the shortest nuclear axis at NEB. Importantly, this temporary prophase centrosome-nucleus configuration is clearly distinct from the mechanism driving metaphase spindle orientation. In this case, external cues [17, 20] activate cortical force generators [16, 18, 19] and transmit pulling forces to astral microtubules [9, 20–22], to ensure robust centrosome positioning. How does cell rounding cooperate with NE Dynein to ensure the correct orientation centrosomes-nucleus axis during prophase? Based on our results and previous reports [47, 48], we propose that Dynein-mediated forces generate a rotational torque on the nucleus that results in centrosome motion when nuclear rotation is limited due to cell adhesion. Alternatively, if cell shape is extensively remodeled during mitotic rounding, Dynein-mediated forces generate nuclear rotation, while centrosomes remain stationary [47]. Accordingly, we show that inducing cell rounding is sufficient to increase nuclear displacement (Fig. S5), whereas forcing cell spreading constrains nuclear rotation (Fig.4 and [49]).

It is unlikely that centrosome movement along the NE surface *per se* justifies the preferential centrosomes-nuclear axis observed at NEB, which argues for an intrinsic nuclear cue in this process. Our results indicate that prophase mitotic chromosome condensation could play such a role, by creating a suitable mechanical environment on the prophase nucleus and allowing NE Dynein loading. During prophase, mitotic chromosomes condense and Lamin A is released into the nucleoplasm [50, 51]. These events likely change the mechanical properties of the nucleus, which in interphase are known to depend on chromosome condensation and the nuclear lamina [34, 36]. How could the mechanical properties of the prophase nucleus regulate Dynein loading? Connection of the nucleus to the cytoskeleton occurs through the LINC complex, composed of KASH proteins on the outer nuclear membrane and SUN proteins on the inner nuclear membrane [52], which ensure nucleo-cytoplasmic mechanotransduction [32, 38]. Importantly, they regulate Dynein loading on the NE [14, 15] to control centrosome tethering and nuclear motility [53]. We propose that mitotic chromosome condensation increases intranuclear stiffness, which is transmitted to the NE through SUN proteins. In turn, this allows Dynein to load on the NE by interacting with NudE/NudEL bound to the nuclear pores through Nup133-CENP-F [8]. In support of this model, it was shown that SUN1 can directly interact with the NPC complex [54], removal of SUN1/2 decreases the accumulation of Dynein on the NE [39] and both SUN1 and SUN2 are necessary to couple chromosomes to the NE [55, 56]. In addition, these observations also justify how confinement, either by inducing transient chromatin condensation [37] or imposing mechanical strain on SUN proteins [31], is able to restore NE Dynein localization. Interestingly, this mechanosensitive behavior seems to be restricted to Dynein, since the Dynein adaptors BicD2 and NudE are still present on the NE even after interfering with chromatin condensation (Fig. S7). The exact nature of the mechanosensitive behavior of Dynein remains to be determined.

Chromosome capture during early mitosis was proposed to occur through a “search-and-capture” mechanism [57]. Subsequent work demonstrated that timely spindle assembly could not rely solely on the “search-and-capture” mechanism [58], but depended on the contribution of kinetochore-driven microtubule nucleation [59], kinetochore compaction [60] and chromosome motion [61] and distribution [62]. In addition to these, we propose an additional spindle assembly facilitating mechanism that requires centrosome-nucleus axis reorientation during prophase to ensure efficient chromosome capture. We propose that centrosome positioning on the shortest nuclear axis favors the assembly of a spindle scaffold to ensure maximum exposure of kinetochores to microtubules. In combination with the spatial distribution of chromosomes in a ring configuration [62], this would accelerate spindle assembly, minimizing the probability of generating erroneous attachments. In agreement with this hypothesis, we observed that centrosome mispositioning significantly delayed mitosis, whereas failure to separate centrosomes altogether generated chromosome missegregation events, as previously described [26].

In summary, we propose a model where centrosome positioning during prophase is a process that depends on an internal mechanosensitive signal provided by chromatin condensation, which facilitates the formation of an initial bipolar spindle scaffold to ensure mitotic fidelity.

## Materials and Methods

### Cell lines and transfections

Cell lines were cultured in Dulbecco’s Modified Eagle Medium (DMEM; Life Technologies) supplemented with 10% fetal bovine serum (FBS; Life Technologies) and grown in a 37°C humidified incubator with 5% CO2. HeLa DHC-GFP and HeLa POM121-3xGFP/H2B-mCherry cell lines were a kind gift from Iain Cheeseman and Katharine Ullman, respectively. HeLa cell line expressing histone H2B-GFP/mRFP-α-tubulin was generated in our lab using lentiviral vectors. For this purpose, HEK293T cells at 50%-70% confluence were co-transfected with lentiviral packaging vectors (16.6 µg of Pax2 and 5.6 µg of pMD2) and 22.3 µg of LV-H2B-GFP (a gift from Elaine Fuchs, Addgene plasmid 25999) or pRRL-mRFP-α-tubulin plasmids, using 30 µg of Lipofectamine2000 (Life Technologies). After transfection, the virus-containing supernatant was collected, centrifuged, filtered and stored at –80°C. HeLa parental cells were then transduced with each lentivirus in the presence of polybrene (1:1000) in standard culture media, for 24 h. The lentiviruses were used individually, giving time for cells to recover between transductions. After the second transduction, H2B-GFP/mRFP-α-tubulin double-positive cells were isolated by fluorescence-activated cell sorting (FACS; FACS Aria II). For transient overexpression of pEGFP-KASH (a gift from Christophe Guilluy) or pRK5-Rap1[Q63E] plasmids (a gift from Jean de Gunzburg), cells were transfected with the corresponding plasmid using Lipofectamine2000 (Life Technologies). Briefly, cells at 50%–70% confluence were incubated for 6 h with 5 µl of Lipofectamine 2000 and 0.6 µg/ml of DNA. DNA-lipid complexes were previously diluted in Opti-Minimal Essential Medium (Opti-MEM; Alfagene) and incubated for 30 min before adding to the cells. Prior to and during transfection, cell medium was changed to a reduced serum medium (DMEM supplemented with 5% FBS). Cells were analysed 48 h after transfection.

### Micro-patterning

Micro-patterns to control individual cell shape and adhesion pattern were produced as previously described[63]. Briefly, glass coverslips (22 × 22mm No. 1.5, VWR) were activated with plasma (Zepto Plasma System, Diener Electronic) for 1 min and incubated with 0.1 mg/ml of PLL(20)-g[3, 5]-PEG(2) (SuSoS) in 10 mM HEPES at pH 7.4, for 1 h, at RT. After rinsing and air-drying, the coverslips were placed on a synthetic quartz photomask (Delta Mask), previously activated with deep-UV light (PSD-UV, Novascan Technologies) for 5 min. 3 µl of MiliQ water were used to seal each coverslip to the mask. The coverslips were then irradiated through the photomask with the UV lamp for 5 min. Afterwards, coverslips were incubated with 25 μg/ml of fibronectin (Sigma-Aldrich) and 5 μg/ml of Alexa546 or 647-conjugated fibrinogen (Thermo Fisher Scientific) in 100 mM NaHCO3 at pH 8.6, for 1 h, at RT. Cells were seeded at a density of 50.000 cells/coverslip and allowed to spread for ∼10-15h before imaging. Non-attached cells were removed by changing the medium ∼2h-5h after seeding.

### Drug treatments

Pertussis toxin (PTx) was used at 40 nM (Merck). Inhibition of Topoisomerase II was done using 10 µM of ICRF-193 (Merck-Millipore). Inhibition of histone deacetylases (HDACs) was done using 1.5 μM of VPA (Sigma-Aldrich). To interfere with the microtubule cytoskeleton, we used nocodazole (20 nM) (Sigma-Aldrich). To inhibit Eg5, STLC was added at 5 μM. All the drugs used were added to the culture medium 30 min-1h before live-cell imaging or fixation, except ICRF-193 and VPA which were added 8h-16h before the experiments. Control cells were treated with DMSO (Sigma-Aldrich) only.

### RNAi experiments

Cells were transfected with small interfering RNAs (siRNAs) using Lipofectamine RNAi Max (Life Technologies). Specifically, 5μl of Lipofectamine and 20 nM of each siRNA were diluted and incubated in Opti-MEM (Alfagene) for 30 min. The siRNA-lipid complexes were then added to 50%–70% confluence cells cultured, during transfection (6 h), in reduced serum medium (DMEM supplemented with 5% FBS). Commercial ON-TARGETplus siRNAs (Dharmacon) were used for Lamin-A/C (set of 4: 5’-GAAGGAGGGUGACCUGAUA-3’, 5’-UCACAGCACGCACGCACUA-3’, 5’-UGAAAGCGCGCAAUACCAA-3’ and 5’-CGUGUGCGCUCGCUGGAAA-3’), BICD2 (SMARTpool: 5’-AGACGGAGCGCGAACAGAA-3’, 5’-UAAAGAAGGUGAGCGACGU-3’, 5’-GCAAGUACCAUGUGGCUGU-3’ and 5’-GGAAGGUGCUAGAGCUGCA-3’) and ARPC4 (set of 4: 5’-GAACUUCUUUAUCCUUCGA-3’, 5’-UAAACCAUCUGGCUGGAUC-3’, 5’-GAAGAGUUCCUUAAGAAUU-3’ and 5’-GAGAUGAAGCUGUCAGUCA-3’) depletions. For Dynein Heavy Chain (DHC) depletion the following oligos were ordered 5’-GAACUAGACUUGGUUAAUU-3’ and 5’-AAUUAACCAAGUCUAGUUC-3’. For combined NudE+NudEL depletion the following oligos were ordered 5’-GCUUGAAUCAGGCCAUCGA-3’ and 5’-UCGAUGGCCUGAUUCAAGC-3’ for NudE and 5’-GGAUGAAGCAAGAGAUUUA-3’and 5’-UAAAUCUCUUGCUUCAUCC-3’for NudEL. Both commercial ON-TARGETplus non-targeting Pool siRNAs (SMARTpool: 5’-UGGUUUACAUGUCGACUAA-3’, 5’-UGGUUUACAUGUUGUGUGA-3’, 5’-UGGUUUACAUGUUUUCUGA-3’ and 5’-UGGUUUACAUGUUUUCCUA-3’) and mock transfections were used as controls. For all siRNAs used, cells were analysed 72 h after transfection. Protein depletion efficiency was monitored by immunoblotting and phenotypic analysis.

### Time-lapse microscopy

For time-lapse microscopy, 12-24 h before the experiments 1.5×10^5^ cells were seeded on coverslips coated with FBN (25μg/ml; F1141, Sigma) or PLL (25μg/ml; F1141, Sigma). When micro-patterns were used, 5×10^4^ cells were seeded on coverslips coated with FBN (25μg/ml; F1141, Sigma). Prior to each experiment, cell culture medium was changed from DMEM with 10% FBS to Leibovitz’s-L15 medium (Life Technologies) supplemented with 10% FBS and Antibiotic-Antimycotic 100X (AAS; Life Technologies). When SiR-dyes were used, they were added to the culture medium 30min-1h before acquisition (20nM Sir-tubulin or 10nM Sir-DNA; Spirochrome). Live-cell imaging was performed using temperature-controlled Nikon TE2000 microscopes equipped with a modified Yokogawa CSU-X1 spinning-disc head (Yokogawa Electric), an electron multiplying iXon+ DU-897 EM-CCD camera (Andor) and a filter-wheel. Three laser lines were used for excitation at 488, 561 and 647nm. For nuclear pore fluctuation analysis, an oil-immersion 100x 1.4 NA Plan-Apo DIC (Nikon) was used. All the remaining experiments were done with an oil-immersion 60x 1.4 NA Plan-Apo DIC (Nikon). Image acquisition was controlled by NIS Elements AR software. For centrosome tracking 17-21 z-stacks with a 0.5µm separation were collected every 20 sec. For mitotic timing quantifications, 13 z-stacks with a 0.7 µm separation were collected every 2 min. For nuclear envelope fluctuation measurements a single z-stack was collected every 100 msec.

### Quantitative analysis of centrosomes, cell membrane and nucleus membrane

Detailed quantitative analysis of centrosomes location and membranes topology (cell and nucleus) was performed using custom made MATLAB scripts (The MathWorks Inc., USA; R2018a). The image analysis took advantage of the different labeling for centrosomes, cell membrane and nuclear membrane. The scripts were separated into three modules with specific workflows: i) centrosomes tracking, ii) nuclear and cellular membrane reconstruction, and iii) nuclear membrane surface dynamics. Tracking of centrosomes position/trajectories was performed in three-dimensional (3D) space using image stacks with a pixel size of 0.190μm and z-step of 0.7μm. Images were pre-processed using a Laplacian of Gaussian filter with a user-defined kernel size, associated with the centrosome radius in pixels. Image segmentation was performed using Otsu’s method, and morphological operators were used to improve the mask and obtain the centrosomes 3D coordinates. Error correction methods, such as automatic thresholding adjustment or in the limit frame elimination, were implemented to take care of frames where the standard method was unable to uniquely identify 2 centrosomes. For the visualization of the centrosomes trajectories (space and time), the centrosomes coordinates were interpolated using cubic splines. Different metrics, such as the distance between centrosomes (pole-to-pole), were calculated to analyze and characterize the trajectories. Cellular and nuclear membranes were reconstructed in 3D space taking advantage of specific labeling. For each membrane, a mask was produced using Otsu’s method and improved with a sequence of morphological operators (namely image close, dilation and erosion, small objects removal). The orientation axis for the membranes were calculated using principal components analysis (PCA) of a large sample of membrane surface points. This method using PCA was found to be more robust than ellipsoid fitting to the membrane surface (followed by extraction of the axis vectors). From the centrosomes locations and nuclear membrane reconstruction, it was possible to calculate the angle between the centrosomes axis and the nucleus major axis. Quantification of nuclear membrane surface fluctuation was performed in 2D using a single slice with a pixel size of 0.102μm. The coordinates of the pixels in the membrane contour were extracted for each frame by first reducing noise with a median filter (neighborhood of 3×3 pixels) followed by object segmentation. The segmentation used a statistical threshold (median + standard deviation), and was improved with small objects removal and closure morphological operations. A reference membrane contour for the nucleus, obtained from the median intensity projection of all frames, was used as “baseline” for the fluctuation analysis. The (Euclidean) coordinates of the nuclear membrane pixels for each frame were converted to polar coordinates, and fluctuations were calculated as the difference in the radial component to the reference contour. The center of the polar coordinates was defined as the centroid of the reference membrane contour. The polar coordinates allowed the decomposition of the fluctuations normal to the nuclear contour (captured in the radial coordinate). The analysis was limited to 60° angular apertures centered on the membranes main axis, as to minimize the error in the radial component. Different methods were designed to explore, analyze and visualize the radial components of the membrane contour. In these methods, the membrane radial fluctuations were characterized using statistics such as maximal amplitudes or standard deviation of the radial component. Radial displacement maps were produced as the radial shift of each point in the membrane with respect to the reference (median) membrane contour. Nuclear irregularity index was calculated as described previously [35].

### Preparation of micropatterned hydrogels with nanobeads

Firstly, 32mm coverslips were plasma cleaned for 30 sec and then incubated with a drop of PLL-PEG 0.1 mg/mL in HEPES 10 mM ph7.4 for 30 min at RT as described previously [64]. Coverslips are then put upright to let the excess PLL-PEG run off and placed on a line or circle shape quartz photomask (Toppan) on a 3μl drop of MilliQ water. The coverslips on the photomask are then exposed to deep-UV for 5min. Then, coverslips are detached from the photomask and incubated with 20μg/ml fibronectin (Sigma) and 20μg/ml Alexa546-conjugated fibrinogen (Invitrogen) in PBS for 30min at RT. To prepare the gels, a 42μl drop of 40KPa mix of Polyacrylamide and bisacrylamide (Sigma) containing 0.1μl carboxylate-modified polystyrene fluorescent beads (Invitrogen) is placed onto the fibronectin coated coverslips and then covered with a second coverslip, pretreated with Bind-silane solution (100% ethanol solution containing 18.5μl Bind Silane; GE Healthcare Life Science) and 161μl 10% acetic acid (Sigma) for 5 min. Gels are polymerized for 30 min and finally the gel is retrieved with the silanized coverslip. Fibronectin proteins are trapped within the acrylamide mesh. Gels are stored in PBS at 4°C.

### Traction force microscopy (TFM) imaging and analyses

For TFM live-cell imaging, rectangle micropatterned coverslips are mounted in dedicated chambers and supplemented with L-15/10% FBS medium. A Leica SP8 confocal microscope was used to acquire the images using a 40X objective (oil immersion, numerical aperture 1.3) with a temperature control chamber set at 37°C. Cells were imaged every 3min. 488 nm and 533 nm lasers were used in sequential scanning mode. All the laser parameters and imaging setups are controlled through the LAS X system. Cellular traction forces were calculated using a method previously described [65]. Briefly, at each time point, the image of the fluorescent beads embedded in the substrate was compared to a reference image corresponding to a relaxed substrate and taken after washing away the cells. After correcting for experimental drift, the displacement field was obtained by a two-step process consisting of cross-correlation on 9.6μm sub-images followed by particle tracking to improve the spatial resolution. The final displacement field was interpolated to a regular grid with 1.2μm spacing. Traction stress reconstruction was performed with the assumption that the substrate is a linear elastic half-space using Fourier transform traction cytometry (FTTC) and zeroth order regularization. The stress map was defined on the same 1.2μm-period grid. From this stress map and the cell mask, we checked that the out of equilibrium force is less than 10% of the sum of forces magnitude, as a quality criterion for all cells and time points. The contractile energy, which is the mechanical energy transferred from the cell to the substrate, was computed from the traction map by integrating the scalar product of the displacement and stress vectors over the cell surface. To determine the principal direction of contraction of each cell, we calculated and diagonalized the first moment tensor of the stress. The eigenvector corresponding to the larger eigenvalue gives the direction of the main force dipole. The degree of force polarization is obtained by comparing both eigenvalues. All the calculations are performed in Matlab (The MathWorks Inc., USA; R2018a).

### Cell confinement setup

For confinement experiments, a dynamic cell confiner was prepared as described previously [66], using a custom-designed layout to fit a 35mm dish. Briefly, a suction cup was made in a polydimethylsiloxane (PDMS, RTV615, GE) mixture (10/1 w/w PDMS A/crosslinker B) in a custom-made mold and baked on an 80°C hot plate for 1h before unmolding. The confining structure on the glass slide was made in PDMS from molds fabricated by standard photolithography. Briefly, an epoxy mold was used with a regular holes array (diameter: 440 μm, 1 mm spacing). A drop of PDMS mixture (8/1 w/w PDMS A/crosslinker B) was poured into the mold. Then, a 10 mm standard microscope coverslip, freshly activated for 2 min in a plasma chamber (Diener Electronics, Germany), was pressed on a PDMS drop to get a residual PDMS layer of minimal thickness. After baking at 95°C on a hot plate for 15 min, excess PDMS was removed. To peel off the glass slide with PDMS pillars, a drop of isopropanol was poured on the slide. Finally, the slide was gently raised by inserting a razor blade between the slide and the mold, allowing the confining glass slides bound to the PDMS structures to be lifted away.

### Immunofluorescence

Cells were fixed with 4% PFA in Cytoskeleton Buffer (274 mM NaCl, 2.2mM Na_2_HPO_4_, 10mM KCL, 0.8 mM KH_2_PO_4_, 4 mM EDTA, 4 mM MgCl_2_, 10 mM Pipes, 10 nM Glucose, pH 6.1) and subsequently permeabilized with 5% Triton X-100 (Sigma-Aldrich) in 1x PBS for 5min. After washing in 10% Triton X-100, cells were blocked with 10% FBS in 10% Triton X-100 in PBS for 30min. All the primary antibodies were diluted in blocking solution and incubated for 1h at room temperature. After this incubation the cells were washed with 10% Triton X-100 in 1x PBS and incubated with the respectively secondary antibody for 1h at room temperature. The secondary antibodies were diluted in blocking solution. DNA was stained with DAPI, which was added to the secondary antibodies solution (1ug/ml, Sigma-Aldrich). After incubation with the secondary antibodies and DAPI the coverslips were washed with 10% Triton X-100 in 1x PBS and sealed on glass slides mounted with 20mM Tris pH8, 0.5 N-propyl gallate and 90% glycerol. The following primary antibodies were used: mouse anti-Lamin A+C (1:500, Abcam), rabbit anti-Lamin B1 (1:500, Abcam), rat anti-alpha Tubulin (1:500 Bio-Rad), rabbit anti-NudE/NudEL antibody (1:500, gift from Richard Vallee), rabbit anti-SUN1 (1:1000, Sigma-Aldrich), rabbit anti-SUN2 (1:1000, Sigma-Aldrich). Alexa Fluor 488, 568 and 647 (1:2000, Invitrogen) were used as secondary antibodies. Where indicated, SiR-actin and SiR-DNA were used at a concentration of 20nM (Spirochrome). Images were acquired using an AxioImager Z1 (63x, Plan oil differential interference contract objective lens, 1.4 NA; all from Carl Zeiss) which is coupled with a CCD camera (ORCA-R2; Hamamatsu Photonics) using the Zen software (Carl Zeiss).

### Quantification of DHC-GFP fluorescence intensity

To quantify fluorescence intensity in immunofluorescence samples, z-stacks containing the entire cell were collected and sum-projected using ImageJ. To identify the nuclear envelope, Lamin B1, SUN1/2 or Lamin A immunostaining were used as a guide. Subsequently, three linescans were done on the nuclear envelope regions corresponding to the long and short nuclear axes, respectively. The mean gray value of each linescan was measured and averaged for each individual cell.

### Western Blotting

HeLa cell extracts were collected after trypsinization and centrifuged at 1200 rpm for 5min, washed and resuspended in 30-50μL of Lysis Buffer (NP-40, 20 nM HEPES/KOH pH 7.9; 1 mM EDTA pH 8; 1 mM EGTA; 150 mM NaCl; 0.5% NP40; 10% glycerol, 1:50 protease inhibitor; 1:100 Phenylmethylsulfonyl fluoride). The samples were then flash frozen in liquid nitrogen and kept on ice for 30min. After centrifugation at 14000 rpm for 8min at 4° C the supernatant was collected and protein concentration determined by the Bradford protein assay (Bio-Rad). The proteins were run on 7%-15% SDS-PAGE gels (50μg per lane) and transferred, using the iBlot Gel Transfer Device (Thermo Scientific), to a nitrocellulose Hybond-C membrane. With the exception of DHC RNAi samples, all others were transferred using a wet blot apparatus for 3h at 70V, with constant amperage. Afterwards the membranes were blocked with 5% Milk in TBS with 0.1% Tween-20 (TBS-T) for 1h at room temperature. The primary antibodies used were: mouse anti-Nde1 (1:500, Abnova), mouse anti-Lamin A+C (1:500, Abcam), mouse anti-LaminA (1:100, Abnova), rabbit anti-ARPC4 (1:500, Bethyl Laboratories), rabbit anti-BICD2 (1:250, Atlas Antibodies), mouse anti-Dynein intermediate chain 74.1 (1:1000, Merck), rat anti-alpha tubulin (1:1000, Bio-Rad). All primary antibodies were incubated overnight at 4°C with shaking. After three washes in TBS-T the membranes were incubated with the secondary antibody for 1h at room temperature. The secondary antibodies used were anti-mouse-HRP and anti-rabbit-HRP at 1:5000. After several washes with TBS-T, the detection was performed with Clarity Western ECL Substrate (Bio-Rad).

### Statistical analysis and data presentation

Each experiment was repeated independently at least three times and sample sizes are defined in each figure legend. We used three to six independent experiments or biologically independent samples for statistical analysis. For knockdown experiments, the knockdown efficiency of each experiment was measured by quantifying immunoblots. When data are represented as box-whisker plots, the box size represents 75% of the population and the line inside the box represents the median of the sample. The size of the bars (whiskers) represents the maximum (in the upper quartile) and the minimum (in the lower quartile) values. Statistical analysis for multiple group comparison was performed using a parametric one-way analysis of variance (ANOVA) when the samples had a normal distribution. When the sample did not have a normal distribution, multiple group comparison was done using a nonparametric ANOVA (Kruskal-Wallis). All pairwise multiple comparisons were subsequently analyzed using either post-hoc Student-Newman-Keuls (parametric) or Dunn’s (nonparametric) tests. When comparing only two experimental groups, a parametric t test was used when the sample had a normal distribution, or a nonparametric Mann-Whitney test was used for samples without normal distribution. Distribution normalities were assessed using the Kolmogorov– Smirnov test. No power calculations were used. All statistical analyses were performed using SigmaStat 3.5 (Systat Software, Inc.).

## Acknowledgments

The authors thank Iain Cheeseman for the HeLa DHC-GFP cell line, Katharine Ullman for the HeLa POM121-3xGFP/H2B-mCherry cell line, Christophe Guilluy for providing the pEGFP-KASH construct, Jean de Gunzburg for the pRK5-Rap1[Q63E] construct, Elaine Fuchs for the LV-H2B-GFP vector and Richard Vallee for the Nde antibody. We thank all members of the CID lab for discussions and suggestions. The authors thank Edgar Gomes, Tiago Dantas, Reto Gassman, Bernardo Orr and António Pereira for critical reading of the manuscript. This work was funded by grants from FEDER – Fundo Europeu de Desenvolvimento Regional funds through the COMPETE 2020 – Operacional Programme for Competitiveness and Internationalization (POCI), Portugal 2020, and by Portuguese funds through FCT – Fundação para a Ciência e a Tecnologia/Ministério da Ciência, Tecnologia e Ensino Superior in the framework of the project PTDC/BEX-BCM/1758/2014 (POCI-01–0145-FEDER-016589). This work was also partially supported by a grant PHC-Pessoa Campus France/Fundação para a Ciência e Tecnologia. V.N. is supported by grant PD/BD/135545/2018 from the BiotechHealth FCT-funded PhD program. M.D. is supported by grant PD/BD/135548/2018 from the BiotechHealth FCT-funded PhD program. Work in the laboratory of H.M. is funded by the European Research Council (ERC) under the European Union’s Horizon 2020 research and innovation program (grant agreement No 681443).

## Author contributions

V.N., M.D and J.G.F. designed and performed experiments. E.V., I.W. and M.B. performed and analyzed TFM experiments. N.C. was involved in setting up the micropatterning technique. P.A. and D.O. developed all MATLAB computational tools. The manuscript was written primarily by J.G.F. and V.N, with significant input by M.P. and H.M. J.G.F. and H.M obtained funding and provided resources.

---

**Figure S1.**
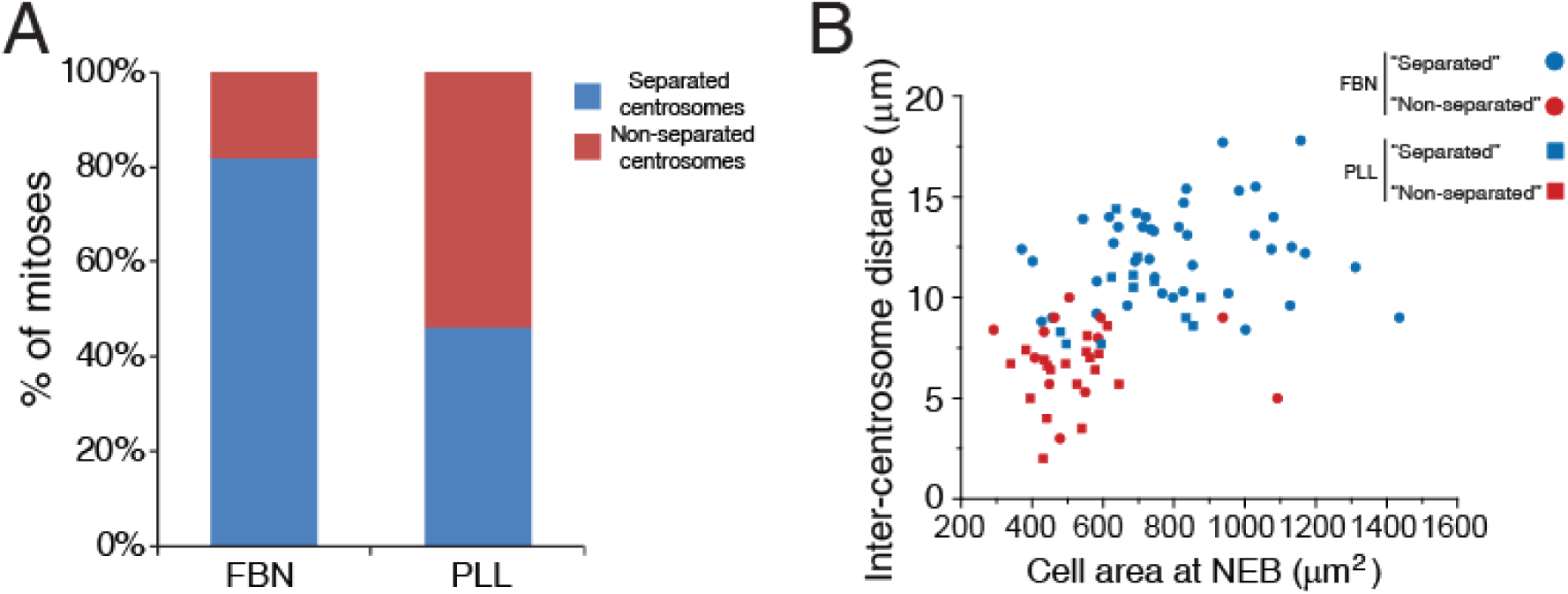
Effect of substrate coating on centrosome separation efficiency, related to Figure 1. (A) HeLa cells expressing H2B-GFP/mRFP-alpha-tubulin were seeded either on FBN (n=59) or PLL (n=41) and imaged to determine whether they separated (“Separated”) or not (“Non-separated”) their centrosomes during mitotic entry. (B) Correlation between inter-centrosome distance (μm) and cell area at NEB (μm^2^) for cells seeded in FBN or PLL, taking into account their centrosome separation pathway. Cells with decreased area have lower inter-centrosome distances.

**Figure S2.**
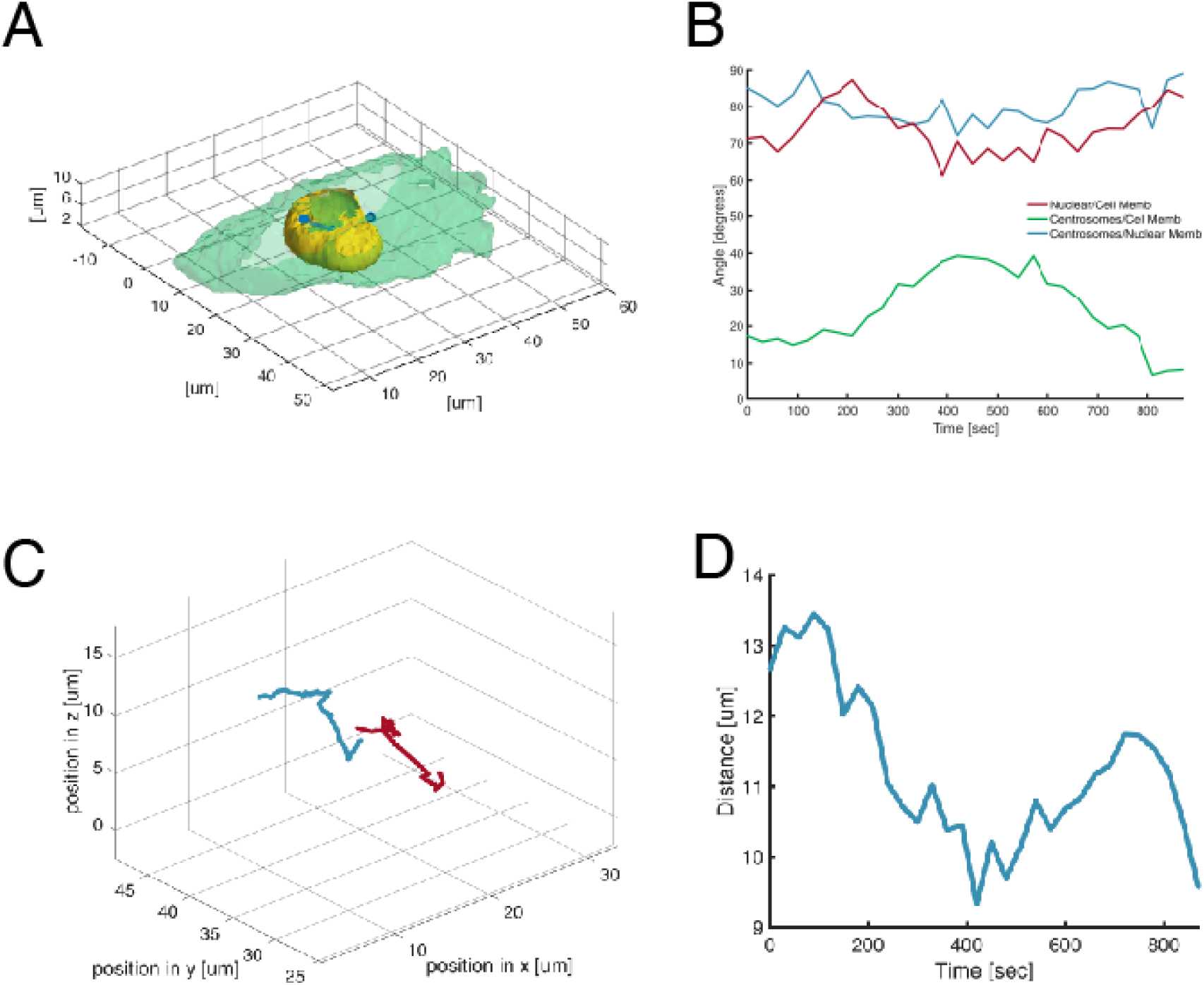
Computational analysis of centrosomes, cell shape and nuclear shape, related to Figure 1. (A) Representative reconstruction of HeLa cell seeded on a line micropattern, showing centrosomes (blue), nucleus (yellow) and cell membrane (green). Centrosomes are connected by a vector that runs through the nucleus centroid. (B) Plot quantifying angles between centrosomes/nucleus long axis (green), centrosomes/cell membrane (blue) and cell membrane/nucleus (red) over time. (C) Representative 4D reconstruction of centrosomes trajectories. (D) Representative plot showing pole-to-pole distance over time.

**Figure S3.**
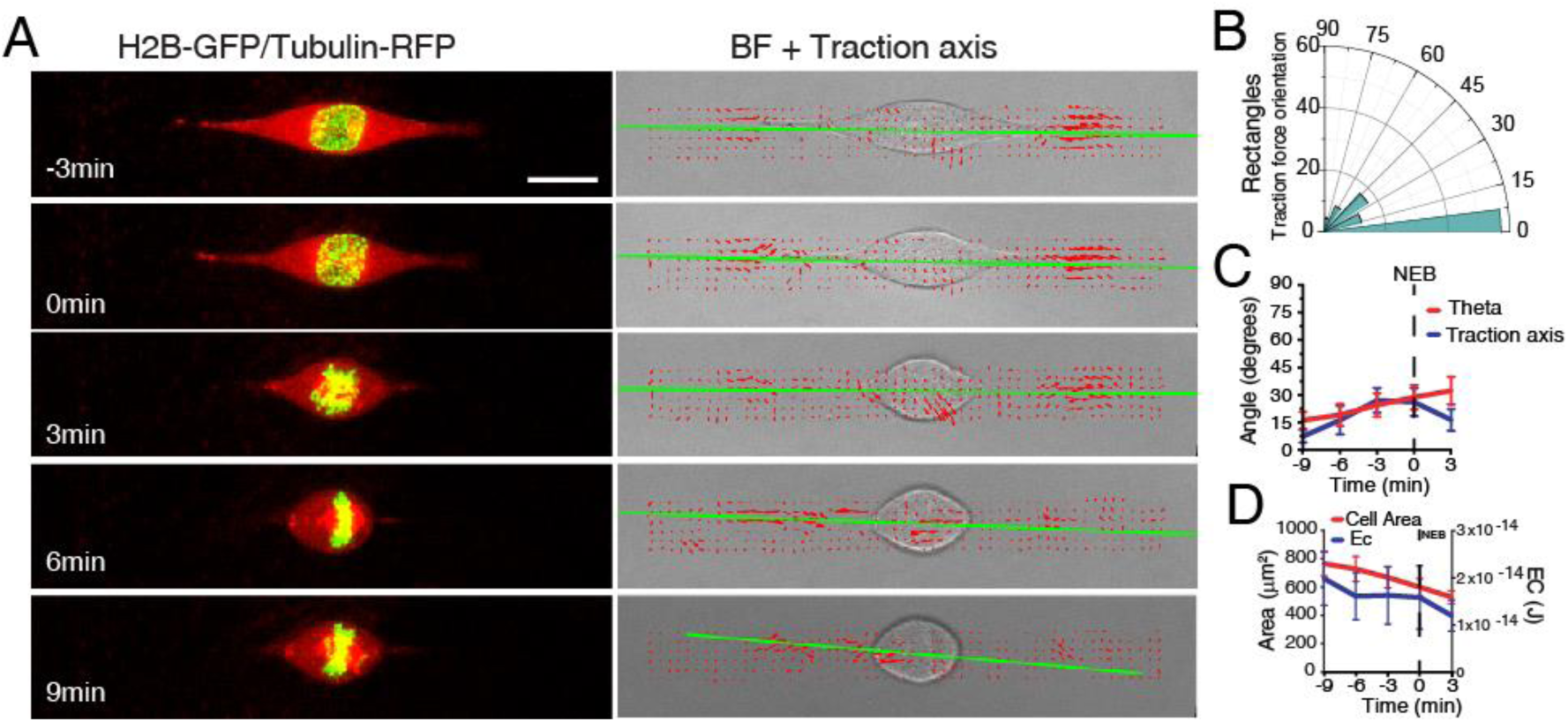
Cell rounding changes the forces exerted by cells on the substrate, related to Figure 4. (A) Frames from a movie of a HeLa cell expressing H2B-GFP/alpha-tubulin-RFP (left panel) seeded on a PAA hydrogel with a rectangle micropattern. Right panel corresponds to brightfield (BF) image overlaid with the corresponding traction force map (red arrows). Green line corresponds to the main direction of the force dipole. Time is in min. NEB = 0min. (B) Traction force orientation for cells on rectangles (n=14). (C) Correlation between centrosome orientation axis (theta; red) and traction axis (blue). Lines corresponds to average values. Error bars correspond to SEM. (D) Correlation between contractile energy (EC; blue) and cell area (red) for cells seeded on rectangles. Lines corresponds to average values. Error bars correspond to SEM.

**Figure S4.**
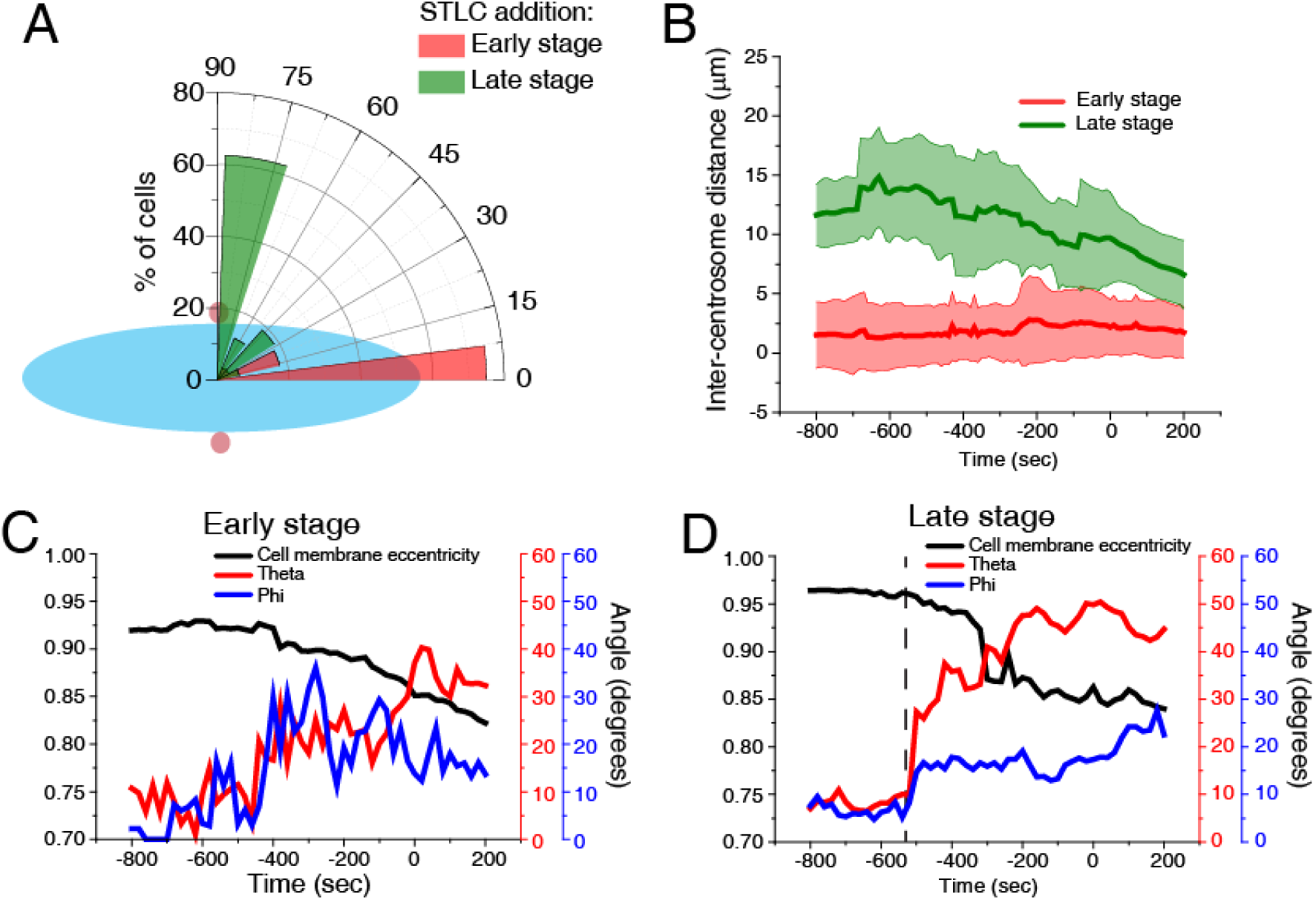
Eg5 is required for early centrosome separation but not centrosome positioning, related to Figure 3. (A) Polar plot quantifying centrosome positioning (red circles) relative to the longest nuclear axis (blue ellipse) at NEB for cells treated with STLC when centrosomes were already on opposite sides of the nucleus (Late stage; green; n=12) or when centrosomes were still not completely separated (Early stage; red; n=30). (B) Inter-centrosome distance for cells treated with STLC in either the Early stage (red) or the Late stage (green) of centrosome separation. Note how inter-centrosome distance only decreases in Late stage cells when cell rounding begins (∼550sec before NEB). Lines correspond to averages and shaded areas correspond to SD. Correlation between the average theta values (red; corresponding to centrosomes xy orientation), average phi values (blue; corresponding to centrosomes z orientation) and cell membrane eccentricity (black) for cells in early (C) or late (D) stage of centrosome separation. The long axis of the micropattern was oriented horizontally and defined as a reference point, corresponding to a value of zero for theta and phi.

**Figure S5.**
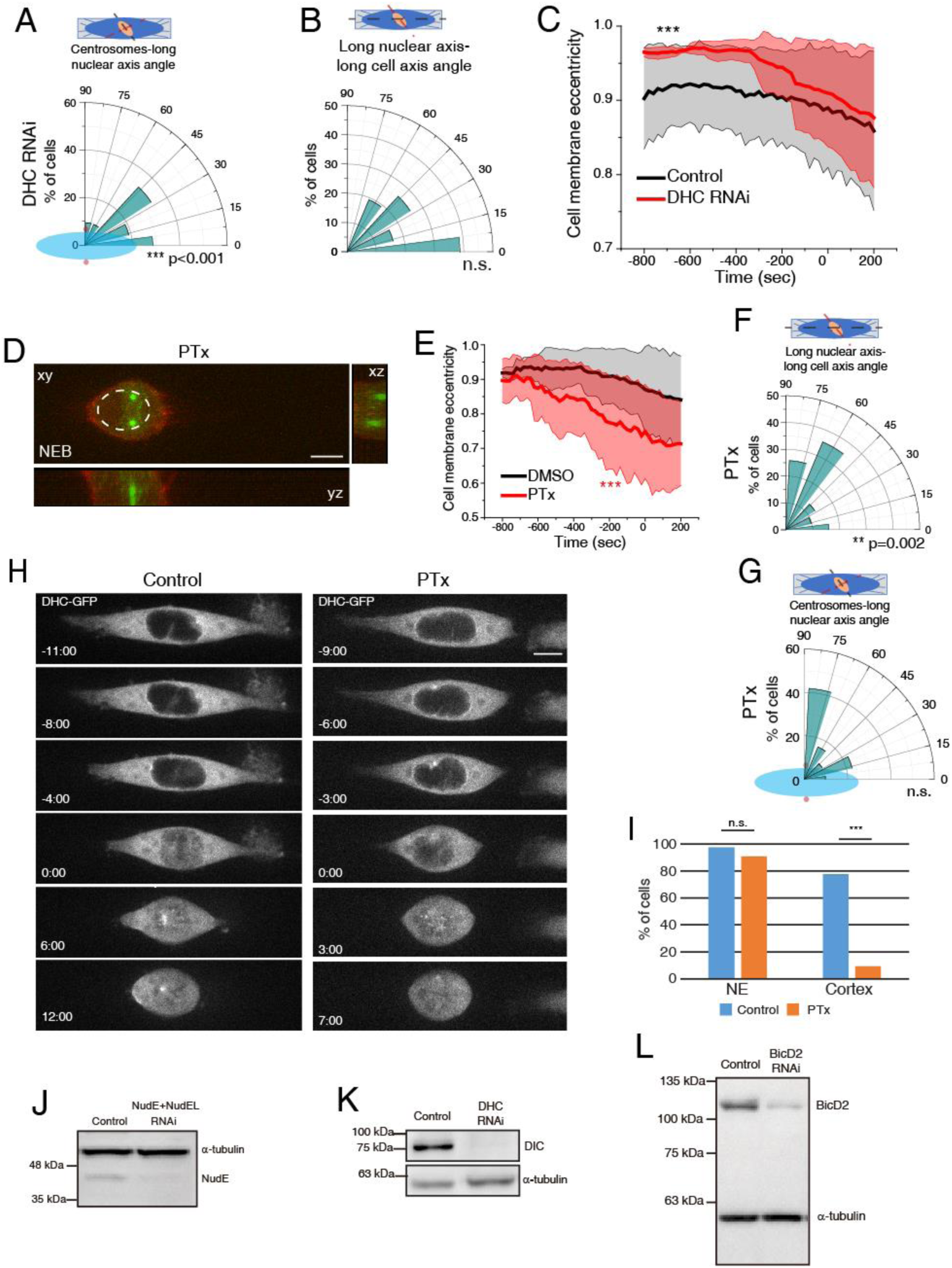
Nuclear envelope Dynein is required for centrosome positioning, related to Figure 4. Polar plots for DHC RNAi-treated cells (n=21) quantifying centrosome positioning (red circles) relative to the longest nuclear (A; *** p<0.01) or alignment of the long nuclear axis with the long cell axis (B) at NEB. (C) Cell membrane eccentricity for controls and DHC RNAi-treated cells. Line correspond to averages and shaded areas correspond to SD (***p<0.001). (D) Representative image of HeLa cell expressing EB3-GFP/Lifeact-mCherry treated with PTx (n=31) at NEB. (E) Cell membrane eccentricity for controls and PTx-treated cells. Line correspond to averages and shaded areas correspond to SD (***p<0.001). Polar plots for PTx-treated cells quantifying alignment of the long nuclear axis with the long cell axis (F; **p=0.002) or centrosome positioning (red circles) relative to the longest nuclear (G) at NEB. (H) Representative images of HeLa DHC-GFP control cells (left panel) or treated with PTx (right panel) during mitotic entry. (I) Quantification of DHC accumulation on the NE and cell cortex. Note how PTx specifically abolishes DHC accumulation on the cell cortex (*** p<0.001). Representative immunoblots to confirm depletion efficiency by RNAi of NudE+NudEL (J), DHC (K) and BicD2 (L).

**Figure S6.**
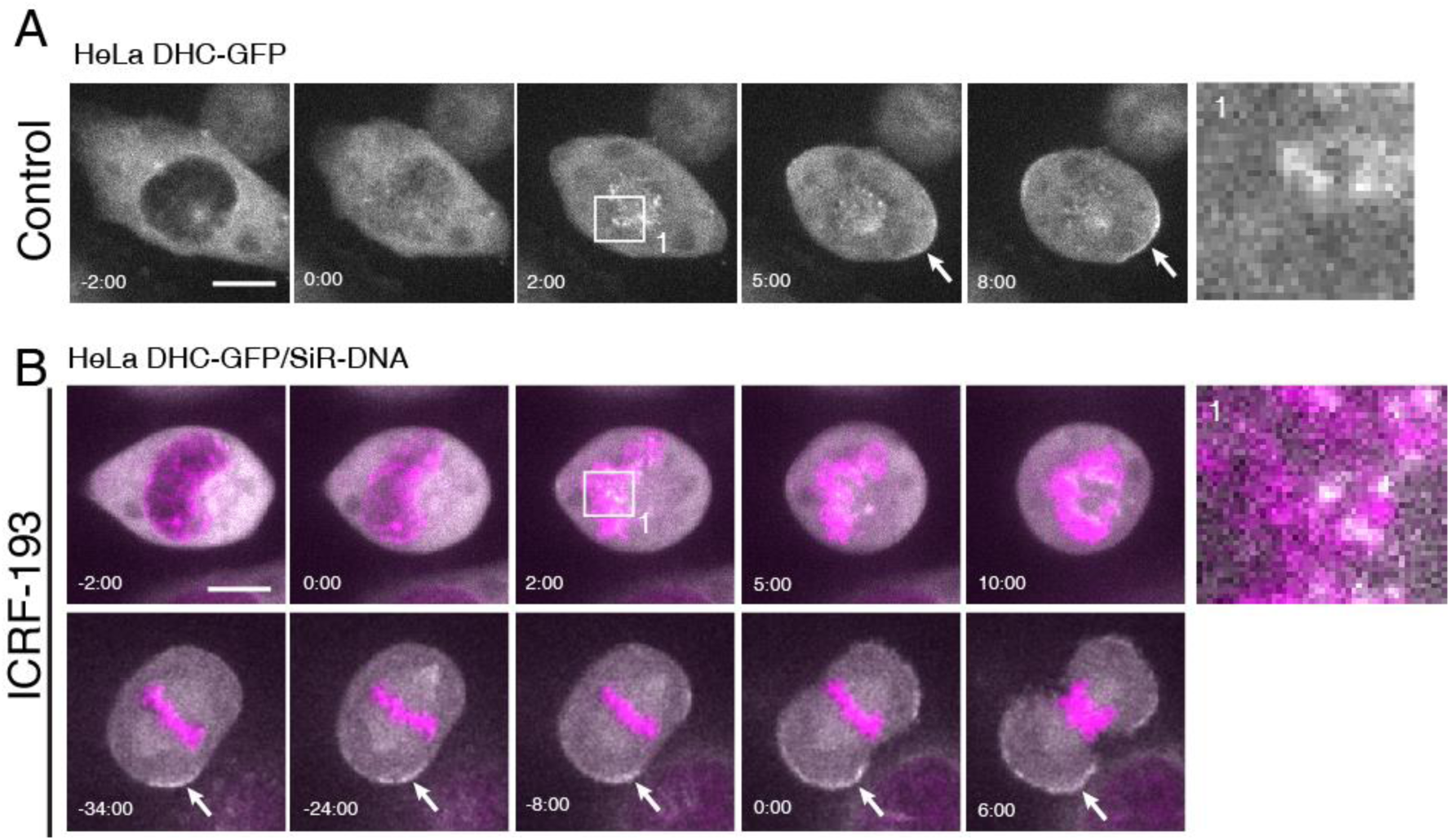
Dynein localizes to kinetochores and cell cortex following treatment with ICRF-193, related to Figure 6. (A) Representative movie of HeLa cell expressing DHC-GFP during mitotic entry, highlighting kinetochore (top panel, white box; 1) and cortical (white arrow) Dynein localization. Time is in min:sec. Time zero corresponds to NEB. (B) Representative movies of HeLa cells labelled with DHC-GFP and SiR-DNA, treated with ICRF-193, highlighting kinetochore (top panel, white box; 2) and cortical (bottom panel) Dynein localization. Time in in min:sec. Time zero corresponds to NEB (top panel) and anaphase onset (bottom panel). Scale bars, 10μm.

**Figure S7.**
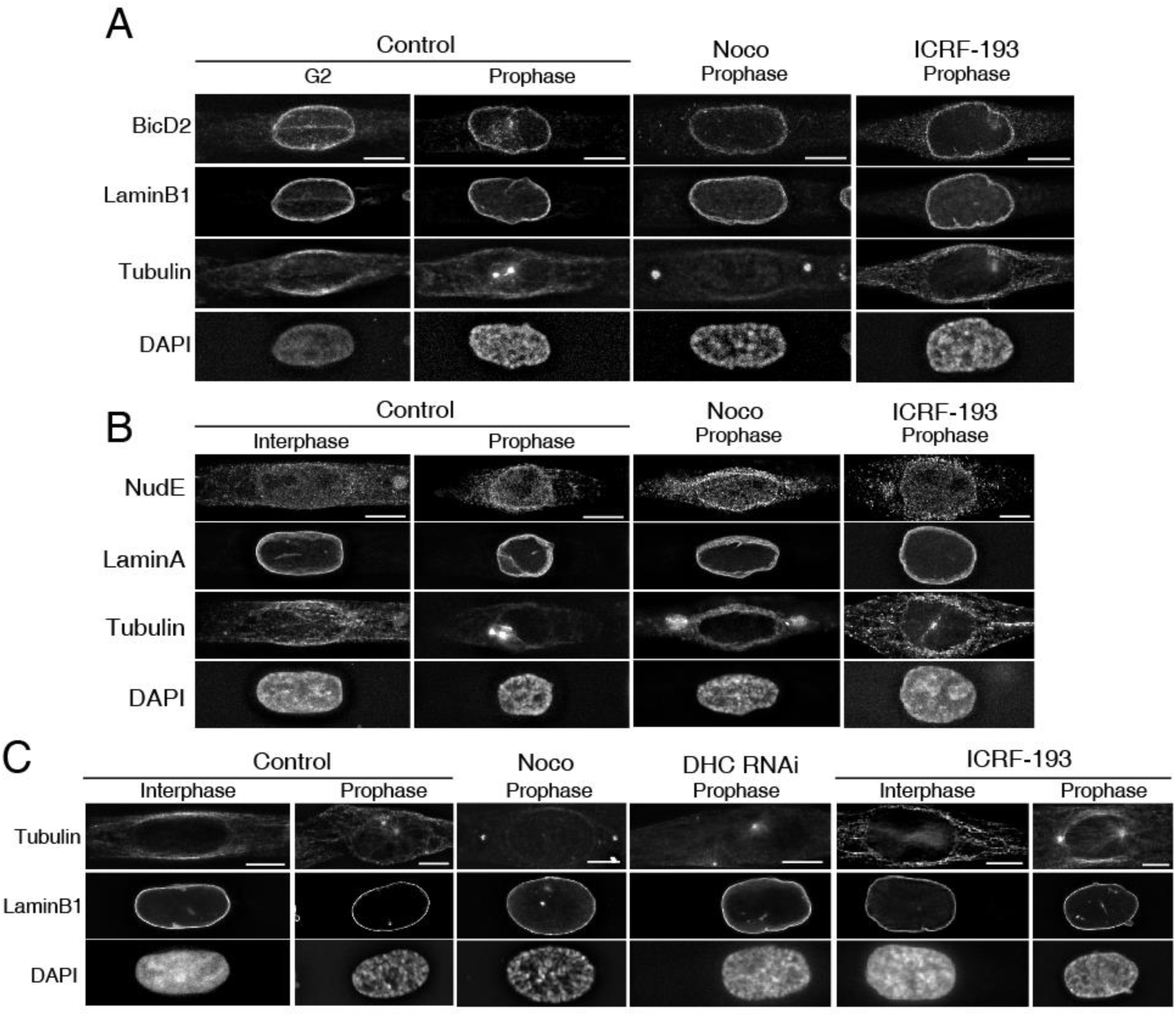
Immunofluorescence analysis of BicD2, NudE, Lamin A and Lamin B1, related to Figure 6. (A) Representative images of immunofluorescence analysis of BicD2, Lamin B1, alpha-tubulin and DAPI for controls (n=31), Noco (n=31) and ICRF-193 (n=30). (B) Representative images of immunofluorescence analysis of NudE, LaminA, alpha-tubulin and DAPI for controls, Noco and ICRF-193. (C) Representative images of immunofluorescence analysis of Lamin B1, alpha-tubulin and DAPI for controls (n=32), Noco (n=32), DHC RNAi (n=29) and ICRF-193 (n=30). All images correspond to deconvolved maximal intensity projections. For quantification purposes, sum-projected images were used. Scale bar, 10μm. All experiments were replicated three times.

**Figure S8.**
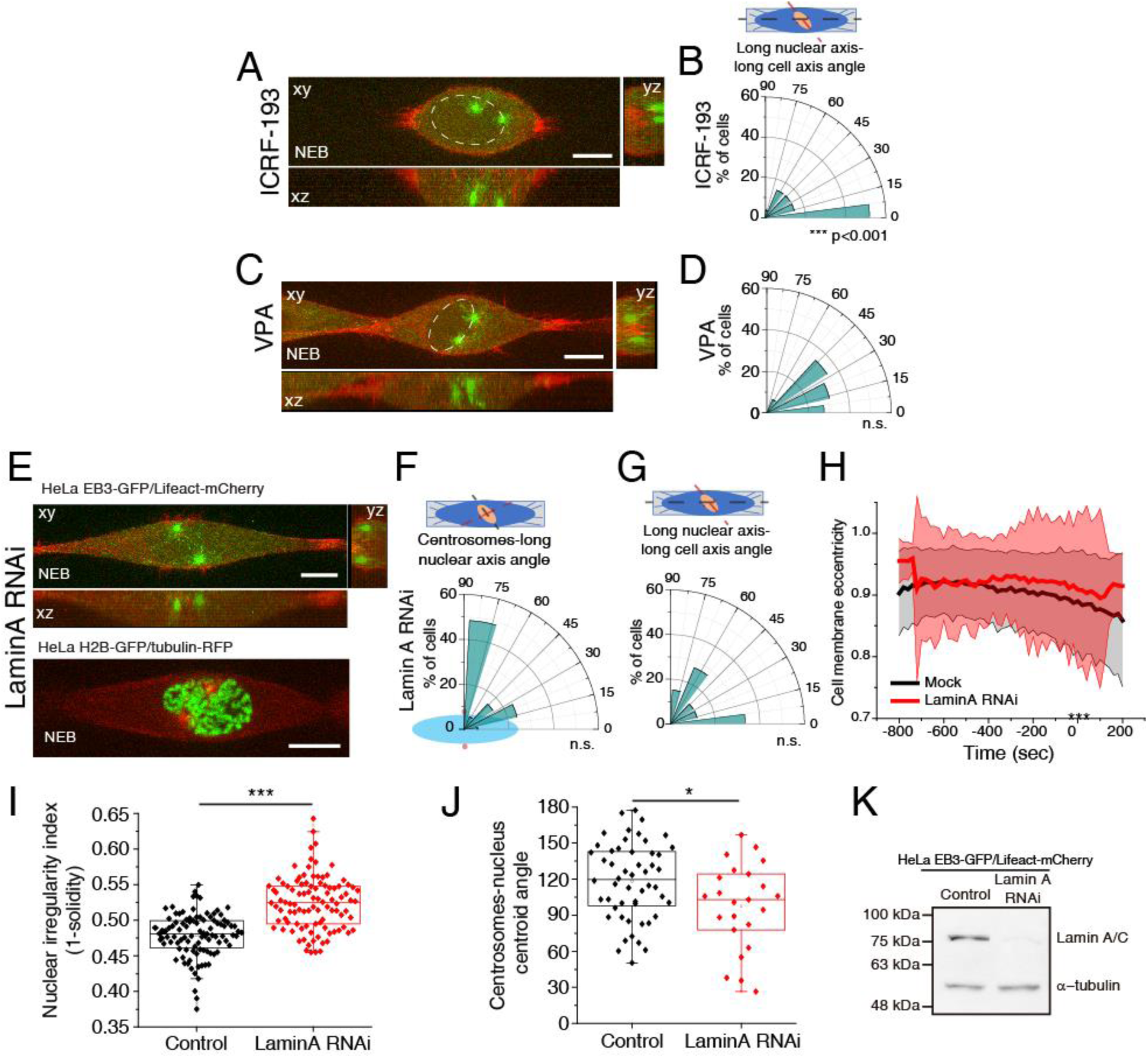
Chromatin condensation is required for centrosome positioning on the shortest nuclear axis, related to Figure 6. (A) Representative time-frame from movie of HeLa cells expressing EB3-GFP/Lifeact-mCherry at NEB, treated with ICRF-193 (n=27). (B) Polar plot for ICRF-193-treated cells quantifying alignment of the long nuclear axis with the long cell axis at NEB. (C) Representative time-frame from movie of HeLa cells expressing EB3-GFP/Lifeact-mCherry at NEB, treated with VPA (n=29). (D) Polar plot for VPA-treated cells quantifying alignment of the long nuclear axis with the long cell axis at NEB. (E) Representative time-frame of cells depleted of Lamin A by RNAi, expressing EB3-GFP/Lifeact-mCherry (n=35; top panel) or H2B-GFP/tubulin-RFP (n=34; bottom panel) at NEB. Note how the nucleus is deformed between the two centrosomes. Polar plots for Lamin A RNAi cells quantifying centrosome positioning (red circles) relative to the longest nuclear axis (F) or alignment of the long nuclear axis with the long cell axis (G) at NEB. (H) Cell membrane eccentricity for controls and LaminA RNAi cells. Line correspond to averages and shaded areas correspond to SD (***p<0.001). (I) Quantification of nuclear irregularity index, calculated as 1-solidity of the nucleus for control (n=15) and Lamin A RNAi-treated cells (n=32; p<0.001). (J) Quantification of the deviation of centrosomes from the nuclear centroid for control and Lamin A RNAi-treated cells (p<0.05). (K) Representative immunoblots to confirm depletion efficiency by RNAi of Lamin A.

